# Stable lentiviral-mediated expression of Cytochrome P450 2D6 in HepaRG cells: New means for *in vitro* assessment of xenobiotic biotransformation and cytotoxicity

**DOI:** 10.1101/2025.01.25.634872

**Authors:** Hugo Coppens-Exandier, Franz Mackanga, Thomas Darde, Corentin Bonhomme, Catherine Ribault, Thomas Gicquel, Romain Pelletier, Flyuza Roshchina, Estelle Schaefer, Christophe Chesné, Agnès Jamin, Anne Corlu, Pascal Loyer

## Abstract

Primary cultures of Human Hepatocytes (PHH) are the gold standard to investigate drug-hepatotoxicity *in vitro*, however, large-scale studies using these primary liver cells are not possible because of the shortage in liver biopsies. HepaRG model is often considered as the closest surrogate to PHH for toxicity studies *in vitro*. However, differentiated HepaRG cells express very low levels of the cytochrome P450 2D6 (CYP2D6) protein, which is essential for the biotransformation of nearly 25% of drugs on the market. To overcome this limitation, infection of progenitor HepaRG cells were performed using lentiviral particles containing a transgene encoding a single mRNA translated into a polypeptide undergoing proteolytic cleavage via the T2A peptide to produce both CYP2D6 and GFP. Differentiated HepaRG cells transduced with lentivirus stably expressed GFP and catalytically active human CYP2D6 enzyme at levels close to those found in high PHH metabolizers. As expected, CYP2D6 protein was found mostly located in the endoplasmic reticulum. Using the CYP2D6 transgenic HepaRG cells, we showed that tramadol was metabolized in both, N- and O-desmethyl tramadol as observed in human serum in contrast with the production of N-desmethyl tramadol only in parental HepaRG cells via the CYP3A4 catalytic activity. Similarly, after perhexiline (PHX) treatments, higher IC_50_ were found in CYP2D6 expressing HepaRG cells associated to lower mitochondrial damages compared to those found in parental cells for the same PHX concentrations. Gene profiling between parental and transgenic cells demonstrated that the CYP2D6 expressing HepaRG cells had kept their ability to proliferate and differentiate with low impact on the expression of the hepatocyte specific functions. However, we identified a limited set of genes such as NXF3 and TRIM63, which were up-regulated by the mRNA encoded by the lentiviral transgene. Together, these data confirmed that the CYP2D6 transgenic HepaRG cells represent a suitable optimized transgenic model of HepaRG cells to evaluate biotransformation and toxicity of specific compounds metabolized by CYP2D6.

## I INTRODUCTION

In the process of drug design, the pharmaceutical industry needs to evaluate the benefits and toxicity of new compounds developed in accordance with ADME (Absorbance, Distribution, Metabolism, Excretion) principals and governmental regulations. The liver is not only the main organ for the biotransformation of xenobiotics, but it is also a major target for their toxicity (Fernandez-Checa et al., 2021 ; Di Zeo-Sánchez et al., 2022). In the context of both acute and chronic exposures, drug-induced liver injury (DILI) is a critical concern for pharmaceutical industry and regulatory agencies and potential hepatotoxicity needs to be carefully evaluated in preclinical drug safety screens using *in vitro* and *in vivo* models (Vinken et al., 2018 ; Fernandez-Checa et al., 2021 ; Daston et al., 2022 ; Braeuning et al., 2023 ; Quintás et al., 2023; Sanz et al., 2023).

While primary cultures of human hepatocytes (PHH) isolated from liver biopsies are considered as the most physiologically cell models to investigate drug-hepatotoxicity *in vitro* (Guillouzo and Guguen-guillouzo, 2008), the shortage in biopsies that can be dissociated to isolate viable hepatocytes, the large variations in the expression of liver specific functions (Guillouzo and Guguen-guillouzo, 2008) and the short lifespan of primary cultures (Gilot et al., 2005) considerably limit the use of PHH on a large scale.

In contrast, well-characterized human hepatoma cell lines including HepG2 and Hep3B (Aden et al., 1979), Huh7 (Nakabayashi et al., 1984) and HBG (Glaise et al., 1998) that can be expanded indefinitely express a limited set of liver specific functions, with very low levels of phase I and II enzymes catalyzing the biotransformation of xenobiotics. Yet, the guidelines for the *in vitro* assessment of hepatotoxicity recommend the use of metabolically competent hepatocyte-like cells (Fernandez-Checa et al., 2021).

The establishment of the human hepatocellular HepaRG cell line twenty years ago provided a new relevant hepatocyte-like model (Gripon et al., 2002), which is now used worldwide for metabolism and toxicity since hepatocyte-like HepaRG cells express most of the hepatocyte specific functions. The HepaRG cells are bipotent hepatic progenitor cells with unlimited proliferation activity that show the remarkable ability to become quiescent at high cell density and to differentiate either towards hepatocyte or biliary epithelial cell lineages producing a coculture system when fully differentiated (Cerec et al., 2007). Progenitor HepaRG cells are considered as a cell model mimicking the differentiation of hepatoblasts into mature hepatocytes (Bucher et al., 2017, Fekir et al., 2019). The optimal differentiation of HepaRG cells in conventional two-dimension (2D) culture is obtained when cells are plated at low cell density to obtain a confluent cell monolayer after two weeks. Then, HepaRG cells are maintained for two more weeks in a culture medium supplemented with 1.5 to 2% dimethyl sulfoxide (DMSO), which strongly enhances the expression levels of a large set of liver specific functions (Cerec et al., 2007 ; Laurent et al., 2013 ; Dubois-Pot-Schneider et al., 2022). Alternatively, 3D culture models of HepaRG cells have also been developed, which may include different cell types combined to HepaRG cells to generate hepatic organoids/spheroids differentiating even in DMSO-free media to avoid the use of this organic solvent (Rose et al., 2022 ; Bernal et al., 2024 ; Bronsard et al., 2024).

A large body of evidences demonstrated the high plasticity of this cell model. Indeed, while progenitor cells are capable of differentiating, differentiated cells are also able to retro-differentiate when quiescent cells are detached and re-plated at low density in absence of DMSO (Cerec et al., 2007). In this condition, both hepatocyte- and cholangiocyte-like cells rapidly lose their differentiation status to retro-differentiated into progenitor cells (Cerec et al., 2007; Dubois-Pot-Schneider et al., 2014). Interestingly, hepatocyte- and cholangiocyte-like cells can be separated by selective detachment (Cerec et al., 2007 ; Laurent 2013) or sorted by FACS and the progenitor cells obtained through retro-differentiation of these two enriched cell populations can differentiate again in both hepatocyte- and cholangiocyte-like cells (Cerec et al., 2007). These experiments demonstrated that the HepaRG cell line consists in a common progenitor cell population giving rise to the two hepatocyte and cholangiocyte lineages. More recently, it was also reported that hepatocyte-like HepaRG cells are not only able to retro-differentiated into progenitor cells but that they can also acquire stem cell features under culture conditions favoring stemness (Dubois-Pot-Schneider et al., 2014 ; Fekir et al., 2019). The HepaRG-derived stem cells express a subset of genes characterizing poorly differentiated and highly proliferative hepatocellular carcinomas of bad prognosis.

The HepaRG-derived stem cells can give rise to highly differentiated hepatocytes, which express most of the drug transporters (Kanebratt and Andersson, 2008 ; Le Vee et al., 2013) and metabolism enzymes (DME) providing a suitable alternative model to PHH for studying the biotransformation (Aninat et al., 2006 ; Kanebratt and Andersson, 2008), toxicity (Jossé et al. 2008 ; Leite et al., 2016 ; Nelson et al., 2017 ; Vlach et al., 2019 ; Bernal et al., 2023) and genotoxicity of xenobiotics (Jossé et al. 2008 ; Kopp et al., 2020 ; Quesnot et al., 2016 ; Thienpont et al., 2023). Since differentiated hepatocyte-like HepaRG cells are relatively stable during several weeks, repeated treatments with xenobiotics can be performed to model chronic exposures (Jossé et al. 2008 ; Leite et al., 2016 ; Kopp et al., 2020).

Several studies have carefully compared gene profiling (Aninat et al., 2006 ; Kanebratt and Andersson, 2008 ; Rogue et al., 2012) as well as protein expression and catalytic activities of drug transporters and DME between PHH and HepaRG cells (Aninat et al., 2006 ; Kanebratt and Andersson, 2008 ; Quesnot et al., 2018). While hepatocyte-like HepaRG cells express nearly 90% of the liver specific functions found in PHH, several important differences have been reported regarding the xenobiotic metabolism. For instance, multidrug resistance proteins MDR 1 and 3, multidrug resistance associated-proteins MRP 1-3, breast cancer resistance protein BCRP (ABCG2), cytochrome P450 CYP3A4, CYP7A1, CYP2C19, glutathione transferase GSTA1 are expressed at higher levels in HepaRG compared to PHH. In contrast CYP1A2, CYP2E1 and bile salt export pump BSEP are found at much lower levels in HepaRG and CYP2D6 is barely detectable (Aninat et al., 2006 ; Kanebratt and Andersson, 2008 ; Quesnot et al., 2018 ; Vlach et al., 2022).

In this context, we and others have developed procedures of transient or stable nucleic acid transfer in HepaRG cells in order to selectively enhance or repress the expression of one or several genes (Laurent et al., 2010 ; Peyta et al., 2016 ; Demazeau et al., 2017 ; Ueyama et al., 2017 ; Pivert et al., 2019 ; Vlach et al., 2019 ; Okuyama S 2022 ; Vlach et al., 2022) without affecting their ability to differentiate and their overall metabolic capacities. Genetic modifications modulating the expression of proteins involved in xenobiotic metabolism is particularly important in order to use HepaRG cells capable of transforming synthetic compounds into their various metabolites and to provide a diversity of genetic backgrounds as observed in humans.

In the past years, we identified several polycationic liposomes that demonstrated efficient plasmid delivery for transient protein expression in proliferating progenitor HepaRG cells (Laurent et al., 2010 ; Demazeau et al., 2017 ; Vlach et al., 2022). However, plasmid transfection of progenitor cells is of limited interest since protein expression will be too transient to be maintained until the cells are differentiated while cells are fully differentiated. Unfortunately, the same transfection reagents yield to very poor reporter gene expression in quiescent differentiated cells (Laurent et al., 2010 ; Vlach et al., 2022) in agreement with the observations that nanoparticle-mediated transfection requires cell division and nuclear envelop breakdown to allow nuclear translocation of plasmids (Zabner et al., 1995 ; Mortimer et al., 1999 ; Tseng et al., 1999). Recently, we developed a mitogenic culture medium that strongly stimulates the proliferation of differentiated hepatocyte-like HepaRG cells (Vlach et al., 2022). The cells are incubated in the proliferation medium during 24 hours, which induces one to two rounds of cell cycle before being switched back to the regular differentiation medium without growth factors to re-induce the quiescence. We showed that high expression levels of CYP2D6 was obtained when lipofection was performed during the 2 to 3-day proliferation period (Vlach et al., 2022). However, we also observed that some of the hepatocyte-specific functions especially phase I enzymes were transiently decreased for 3 to 4 days when HepaRG cells proliferated indicating that the maximal expression of the CYP2D6 was achieved concomitantly to a significant reduction of some other metabolic pathways (Vlach et al., 2022).

To achieve expression of a protein of interest in differentiated HepaRG cells, the transfection of a plasmid encoding the cDNA of this protein and a selection marker such as an antibiotic can appear as a suitable approach. Based on our experience, plasmid transfection followed by strong selection pressure using antibiotics to eliminate cells that do not stably integrate the transgene into their genome impairs the ability of HepaRG cells to properly proliferate and differentiate over a long period of time. However, using this approach Okuyama and co-workers have recently obtained several clones of HepaRG cells that express different levels of CYP2D6 although they did not characterize either the CYP2D6 catalytic activities nor the expression levels of other DMEs in these transgenic lines (Okuyama et al., 2022). Alternatively, the infection of progenitor or differentiated HepaRG cells with lentiviral particles is a straight forward and very efficient procedure to obtain stable transgenic cells without antibiotic selection or for a short period of time that does not compromise the differentiation (van der Mark et al, 2017. ; Quesnot et al., 2018 ; Pivert et al., 2019).

Among the CYP450 family, CYP2D6 plays a central role in drug metabolism since 20 to 30% of drugs are oxidized by the CYP2D6 enzyme. For example, analgesics such as tramadol and codein, antidepressant or anti-tumoral drugs like tamoxifen are known to be metabolized by CYP2D6. In this study, our main goal was to establish a transgenic HepaRG cell line using lentiviral transduction in order to express high levels of CYP2D6 with a catalytic activity in the same range of those found in high metabolizer PHH. Then, we used two compounds known to be hydroxylated by CYP2D6, tramadol and perhexiline, to evaluate their biotransformation and cytotoxicity, respectively, in both parental and the CYP2D6 expressing HepaRG cells. Gene expression profiling between the parental and the CYP2D6 expressing HepaRG cells was performed to determine the impact of the lentiviral transgene integration and/or the enforced expression of CYP2D6 on the differentiation process of genetically modified cells.

## II MATERIALS AND METHODS

### 2.1. HepaRG^TM^ cell culture

The progenitor parental cells (code HPR101, Biopredic^International^, Saint Grégoire, France) were cultured following well-standardized procedures (Supporting information 1) in William’s E growth culture medium (reference # MIL710, Biopredic^International^) composed of basal hepatic cell medium (# MIL700, Biopredic^International^) supplemented with HepaRG™ Growth Medium Supplement (# ADD710, Biopredic^International^) as previously described (Vlach et al, 2022). After plating, the HepaRG cells actively proliferate during the first week before progressively entering quiescence and committing to hepatocyte or cholangiocyte lineages during the second week post-plating (Cerec et al., 2007 ; Dubois-Pot-Schneider et al., 2022). In order to obtain the highest level of differentiation, confluent quiescent cells were cultured for 2 more weeks in Differentiation Medium (# MIL720, Biopredic^International^) composed of basal hepatic cell medium (# MIL700, Biopredic^International^) supplemented with HepaRG™ differentiation medium Supplement (# ADD720, Biopredic^International^) containing 1.7% DMSO, which strongly potentiates the hepatocyte differentiation (Dubois-Pot-Schneider et al., 2022). At day 30 of culture, culture medium of differentiated HepaRG^TM^ cells was discarded, cells were rinsed with PBS prior detachment by incubation with trypsin (Cerec et al., 2007 ; Laurent et al., 2013). Cell suspensions were centrifuged and pellets were resuspended in cryogenic medium prior freezing and storage in liquid nitrogen. These frozen differentiated cells provide functional hepatocyte-like cells (**Supporting information 1**, https://wepredic.com/en/hepargtm), which can be plated and kept for one week to in appropriate culture medium to obtain the highest levels of specific functions.

Differentiated HepaRG™ cell cryogenic vials (code HPR116, Biopredic^International^) were thawed (at least one week after freezing) and plated in HepaRG™ Thawing/Plating/General Purpose medium (# MIL670; Biopredic^International^) composed of Basal hepatic cell medium (# MIL600 ; Biopredic^International^) supplemented with HepaRG™ Thawing/Plating/General Purpose medium supplement (ADD670). Twenty-four hours after plating, this medium was removed and cells were then cultured in HepaRG™ Maintenance/Metabolism medium (MIL600) supplemented with HepaRG™ Maintenance/Metabolism medium supplement (ADD620) for at least 7 days prior to be used for gene profiling, metabolism and cytotoxicity assays.

To study the hepatocytes and cholangiocytes populations separately, a selective hepatocyte and cholangiocyte detachment is performed. For a 75 cm^2^ flask, 2.5 mL of trypsin are mixed with 2.5 mL of PBS. Then, the flask was incubated at room temperature and the detachment of hepatocytes is monitored using a phase contrast microscope. After 5 to 10 min, when hepatocyte-like cells began to round up, the flask was gently shaken at room temperature and the complete detachment of hepatocytes was continually monitored under microscope. When almost all the hepatocytes were detached, the supernatant was collected and 5mL of warm MIL670 was added to inactivate the trypsin. This supernatant was centrifuged (2 min, 150g) and the hepatocyte enriched fraction was resuspended in medium. The flask was washed twice with PBS and the remaining cholangiocyte colonies were detached by incubating for 5 min with 2mL trypsin at 37°C. Then, 5 mL of MIL670 was added and the cells were collected. This cell suspension was centrifuged and cholangiocyte enriched fraction was resuspended in MIL670 culture medium. Both fractions were counted and seeded at high density (2,5×10^5^ cells/cm^2^). After 24h, the cultures were switched to MIL620 culture medium. The cultures were maintained in MIL620 culture medium before being used for further studies.

### 2.2. Lentiviral transduction and production of stable recombinant CYP2D6-HepaRG^TM^ cells

The lentiviral particles were produced by Flash Therapeutics (Toulouse, France). Shortly, the hCYP2D6wt cDNA sequence (NM_000106.4) was cloned into a lentiviral backbone also encoding for the Green Fluorescent Protein (GFP) and separated by the sequence T2A proteolytic cleavage site (Supplementary data). The lentiviral particles were produced at 10^7^ Pfu/mL. Transduction of HepaRG^TM^ cells were performed 24h after cell seeding of progenitor HepaRG cells with a theorical multiplicity of infection at 1 infectious viral particle per cell. Then, lentivirus containing medium was discarded and the cells were maintained in culture conditions of parental cells up to 14 days post-seeding before being split and replated at low density in order to expand the cell population. A working bank was created after 3 successive passages and vials of CYP2D6 expressing HepaRG cells at day 14 after seeding, prior differentiation, were frozen and stored in liquid nitrogen. The HepaRG cells expressing CYP2D6 were named HPR1LV01 and HPR6LV01 for progenitor and differentiated cells, respectively (**Supporting information 1**).

### 2.3. Invalidation of the CYP2D6-GFP transgene by CRISPR/Cas9 technology

The Gene Knockout kit (CRISPR-Cas9 kit) targeting human CYP2D6 purchased form Synthego (Redwood, CA, USA) contained three different sgRNA (**Supporting Information-Table 1**) and a Cas9 protein batch. sgRNA were resuspended at 30µM in buffer R while Cas9 was ready to use at 20µM. Ribonucleo protein (RNP) were formed at 3 ratios sgRNA:Cas9 (4:1, 6:1, 8:1) in 35µL of the electroporation buffer R for 10 min. Then, 7.5×10^5^ progenitor HepaRG CYP2D6^+^ cells were detached by trypsin, rinsed in PBS and resuspended in 25µL of buffer R. RNP and cells in buffer R were mixed (60µL final) and 10µL of the mix was electroporated at 1700V, 20ms, 1 pulse with the Invitrogen™ Neon™ Transfection System. The operation was repeated five times to obtain approximately 5×10^5^ electroporated cells seeded in MIL710 warm medium.

After one week, the cells which were efficiently transfected lost the GFP fluorescence. HepaRG cells were collected, filtered through a cell strainer (40µm) and negative GFP cells were sorted using a BD FACS Aria cell sorter (cytometry core facility of the Biology and Health Federative research structure Biosit, Rennes, France). This enriched GFP negative population was expanded and used for further experiments. CRISPR-Cas9-invalidation of the CYP2D6-GFP transgene was verified by PCR using Platinum™ SuperFi™ II Green PCR Master Mix with specific primers and by sequencing performed by Synthego using specific primer and using ICE Synthego online software (https://ice.synthego.com)

### 2.4. Co-localization of CYP2D6, mitochondria and endoplasmic reticulum

Co-localization of CYP2D6 with the mitochondria and the endoplasmic reticulum was performed *via* Mitotracker Red CMXRos (M7512, ThermoFischer) stain and immunostaining of Endoplasmic Reticulum Aminopeptidase 2 (ERAP2). Differentiated WT HepaRG and CYP2D6+ HepaRG cells were seeded on Lab-Tek chamber slides (ThermoFischer, 155411pk) for one week. First, some cells were incubated for 30 min with 0.5µM of Mitotracker Red CMXRos. Then, all the cells were fixed with a cold 4% paraformaldehyde solution for 20 min at room temperature and rinsed with 1xDPBS (Gibco, 14190144). Cells were then incubated for 45 min at room temperature in a blocking solution (DPBS, 0.2% BSA, 0.2% saponin, 0.1% sodium azide) and incubated for 1h at room temperature with a combination of primary antibody: anti-CYP2D6 (738667S, Cell Signaling) and Mitotracker Red CMXRos or anti-CYP2D6 and ERAP2 (MAB3830, BioTechne) diluted at 1:250 in the blocking solution. After rinsing, cells were incubated for 45min at room temperature with secondary antibody: anti-rabbit Alexa Fluor 647 (A31573, ThermoFischer) for CYP2D6 and anti-goat Alexa Fluor 568 (A10037, ThermoFischer) for ERAP2, both diluted at 1:250 in the blocking buffer. Finally, HepaRG cells were rinsed and stained with Hoechst 33342 (62249, ThermoFischer) for 15 min. Colocalization of CYP2D6 with mitochondria or endoplasmic reticulum was performed by using JACoP plugin (V2.1.4) from Bolte, S., & Cordelières, F. P. (2006) on ImageJ. It consisted in superimposing 0.3 μm sections from the confocal image Z-stack, taken with a ZEISS LSM 980 Airyscan 2 confocal microscope, and calculate a Pearson’s coefficient between 2 stains.

### 2.5. Flow cytometry

To analyze transfection efficiency, the number of GFP positive cells and the mean of fluorescence were measured by flow cytometry with FITC-A laser. The cells were detached with trypsin, resuspended in William’s E medium containing FCS. The fluorescence intensity of 10,000 cells was analyzed with a Becton Dickinson LSRFortessa™ X-20 (cytometry core facility of the Biology and Health Federative research structure Biosit, Rennes, France). The mean of fluorescence intensity (MFI) was measured on the GFP positive cell population, which reflects the relative GFP expression levels in the cell population. Flow cytometry data were analyzed using FlowLogic software (7.2.1 version, Inivai Technologies, Mentone Victoria, Australia).

### 2.6. Quantification of phase I and II enzyme activities by LC/MS-MS

For the determination of phase I and II enzymes we used two different cocktails depending on the experimental conditions, either maximum velocity (Vmax) cocktail or hepatic clearance (Clint) cocktail.

For the phase I Vmax cocktail, HepaRG™ cells were incubated on MW96 with a mix of phase I probe substrates, i.e. phenacetin (CYP1A2, 200µM), bupropion (CYP2B6, 100µM), dextromethorphan (CYP2D6, 100µM) purchased from Sigma, and midazolam (CYP3A4/5, 50µM, Pharmacopée Européenne). For phase II Vmax cocktail, HepaRG™ cells were incubated on MW96 with paracetamol (1mM, Sigma) for 1h. Paracetamol sulfation and glucuronidation activities were evaluated. The reaction was stopped by addition of ice-cold acetonitrile and supernatants were collected after centrifugation. The samples were stored at - 20°C until analysis.

For the phase I Clint cocktail, HepaRG™ cells were incubated on MW12 at a concentration of 10^6^ cells/mL with a mix of phase I probe substrates (Xenomix™ 1S, #SL41QC03 ; Starlight, Orléans, France), i.e. phenacetin (CYP1A2, 1µM), coumarin (CYP2A6, 1µM), bupropion (CYP2B6, 1µM), amodiaquine (CYP2C8, 1µM), diclofenac (CYP2C9, 1µM), Mephenytoin (CYP2C19, 5µM), dextromethorphan (CYP2D6, 1µM), testosterone (CYP3A4, 5µM) and nifedipine (CYP3A4/5, 1µM). The reactions were stopped by addition of ice-cold acetonitrile after 5, 10, 15, 20, 30, 45 and 60 min and supernatants were collected after centrifugation. The samples were stored at -20°C until analysis.

The day of analysis, 50 µL (Clint) or 10 µL (Vmax) aliquot of supernatant was transferred to a 96-well plate with round bottom containing 50 µL or 90 µL respectively, of an internal standard working solution. The formed metabolites were quantified by liquid chromatography coupled to tandem Mass Spectrometry (LC/MS-MS).

The analysis was performed on an Exion LC AC system (Applied Biosystems/Sciex, USA). Compounds were separated on a Xselect HSS T3 column (2.1×75mm, 2.5µm, Waters, Milford, MA) maintained at 35°C. The mobile phase was composed of 2 mM ammonium formate containing 0.1% formic acid (A) and acetonitrile (B). Detection was operated on a triple quadrupole mass spectrometer, AB Sciex Triple Quad 5500+ (Applied Biosystems/Sciex, USA), equipped with an electrospray ion source in positive mode. Acquisition was performed using the multiple reaction monitoring (MRM). Data processing was performed using Analyst 1.7.2 software.

Vmax was calculated using this equation:

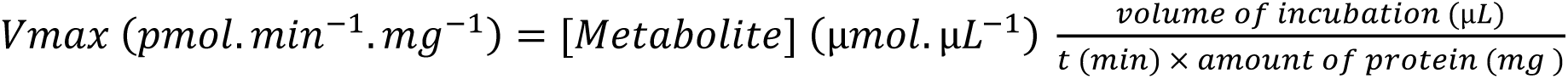

Cl_int_ was calculated using this equation:

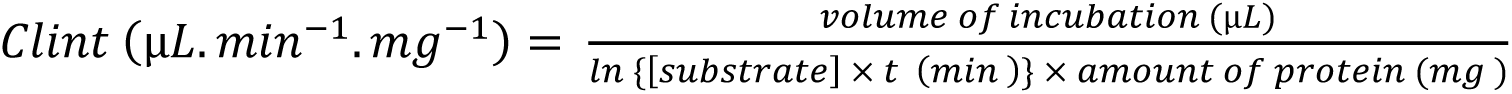

Protein

### 2.7. Induction of CYP450 genes and inhibition of the CYP2D6 catalytic activity

For induction, HPR116 and HPR6LV01 were thawed in MW96 plate at 72.000 cells/well for three days in MIL670 before use. Next, induction experiments were performed in MIL640 (MIL600+ADD640 induction medium) completed with three different inductors (0,1%DMSO) for two days. Omeprazole (CYP1A2), rifampicin (CYP3A4) and phenobarbital (CYP2B6) were diluted at 50µM, 10µM and 1mM respectively in MIL640 warm medium and incubate with the cells for 24h. Medium with inductors was changed the next day for the same medium with inductors for 24h more. After 48h, CYP450 activities were measured with Vmax cocktail as described above.

Quinidine is a potent competitive inhibitor of CYP2D6. To study inhibition of the overexpressed CYP2D6, HPR6LV01 cells are thawed in MW96 plate at 72.000 cells/well for three days in MIL670. Then the medium was changed for MIL620 for four days. After 7 days, cells were incubated with 1 µM or 10 µM of quinidine and CYP2D6 activity was measured with dextromethorphan in Vmax as described above.

### 2.8. Tramadol biotransformation

HPR116 and HPR6LV01 were thawed in MW96 plate at 10^5^ cells/well for three days in MIL670. Then the medium was changed for MIL620 for four days. After 7 days, cells were incubated with 20µM of tramadol diluted in physiological serum (<0,1%) for 48h. Then, supernatants were collected and dissolved in 200μL of water/acetonitrile/formic acid (90/10/0.09, v/v/v). The final extracts were transferred into vials for LC–MS/MS analysis and quantitation as already described (Pelletier et al., 2022) and detailed in **Supporting Information-Table 2**. Results were expressed in µg/L.

### 2.9. BRB-seq library preparation and sequencing

The 3’ Bulk RNA Barcoding and sequencing (BRB-seq) (Alpern et al., 2019) experiments were performed as previously described (Giacosa et al., 2021). Briefly a first step of reverse transcription and template switching reactions were performed using 4 µL total RNA at 2.5 ng/µL. Then cDNAs were purified and Double strand (ds) cDNAs were generated by PCR. The sequencing libraries were built by tagmentation using 50 ng of ds cDNA with the Illumina Nextera XT Kit (Illumina, #FC-131-1024) following the manufacturer’s recommendations. The resulting library was sequenced on a NovaSeq sequencer following Illumina’s instructions by the IntegraGen Company (https://integragen.com/fr/). Image analysis and base calling were performed using RTA 2.7.7 and bcl2fastq 2.17.1.14. Adapter dimer reads were removed using DimerRemover (https://sourceforge.net/projects/dimerremover/).

Briefly the first read contains 16 bases with a higher quality score than 10. The first 6 bp correspond to a unique sample-specific barcode and the following 10 bp to a unique molecular identifier (UMI). The second reads were aligned to the human reference transcriptome from the UCSC website (release hg38, downloaded in August 2020) using BWA version 0.7.4.4 with the parameter “−l 24”. Reads mapping to several positions in the genome were filtered out from the analysis. The pipeline is described in (Giacosa et al., 2021). After quality control and data preprocessing, a gene count matrix was generated by counting the number of unique UMIs associated with each gene in lines for each sample in columns. The UMI matrix was further normalized with the regularized log (rlog) transformation package implemented in the DeSeq2 package.

Statistical filtration analysis was performed using R v4.0.3. A Uniform Manifold Approximation and Projection for Dimension Reduction (UMAP) (https://doi.org/10.48550/arXiv.1802.03426) was performed with the UMAP package implemented in R . Differentially expressed genes were spotted by comparing each condition. Briefly, different filtration steps were applied, including for each comparison: i) median gene expression of all the samples as a background cut-off; ii) a fold-change cut-off of 1.5 (determined using the elbow method) compared to control samples which has been chosen on the base of the curve and iii) a statistical filtration with the linear models for Microarray Data (LIMMA) package and a p-value cut-off of 0.05 adjusted with the Benjamini and Hochberg method. The genes differentially expressed have also been visualized as heatmap representations generated with the R package heatmap.

A Gene Ontology enrichment analysis was performed with the Annotation Mapping Expression and Network (AMEN) suite. A specific annotation term was considered enriched in a gene cluster when the False Discovery Rate (FDR)-adjusted p-value (Fisher’s exact probability) was ≤ 0.05 and the number of genes associated.

### 2.10. Gene expression by RT-qPCR and immunoblotting

After discarding culture medium and washing with PBS, cells were lysed, and total RNA were extracted using Macherey-Nagel NucleoSpin® RNA kit. Following elution, quantification of total RNAs was performed by measuring optical density at 260 nm. Then, reverse transcription was performed using 0,5 µg of total RNAs using the High-capacity cDNA reverse transcription kit (Applied Biosystems). RNA expression levels of specific genes (**Supporting Information-Table 3**) were measured with SYBR Green PCR Master Mix kit (Applied Biosystems) using the QuantStudio^TM^ 7 flex instrument (Applied Biosystems). TATAbox Binding Protein (TBP) genes was used as reference gene.

For immunoblotting, culture medium was discarded, and cells were washed with PBS and lyzed using lysis buffer: Tris 25 mM pH 7.4, 0.1% SDS, NaCl 150 mM, 1%NP-40 and 0.5% sodium deoxycholate supplemented with protease inhibitors (EDTA-free, Roche). Samples were then sonicated, and their total protein contents were quantified using BCA Pierce protein assay (Ref: 23225) in total cell lysates. For immunoblotting, 30 µg of protein was mixed with Laemmli buffer and boiled during 10 min. The samples were next separated on NuPAGE® Novex® Bis-Tris 4–12% gels kit (Invitrogen) and transferred to PVDF membranes (Trans-blot® Turbo™ Transfer System, Biorad) before performing immunoblotting using the following primary antibodies against: GFP (anti GFP goat polyclonal antibody, sc-5384, Santa Cruz Biotechnology), CYP2D6 (anti-CYP2D6, 738667S, Cell Signaling) and HSC70 (anti-HSC, mouse monoclonal antibody, sc-7298, Santa Cruz Biotechnology). Primary antibodies were detected using secondary rabbit or mouse antibodies coupled to horseradish peroxidase (HRP) (Dako, Denmark). Detection of the immune complex was performed using a chemiluminescent HRP substrate (Pierce™ ECL Substrate). The proteins were visualized with the Fusion FX system (Vilber-Lourmat, Eberhardzell Germany).

### 2.11. Cell viability and toxicity assays

Cell viability was evaluated by measuring with the relative intracellular amounts of ATP after treatment with perhexiline (PHX) at 75, 50, 37.5, 25, 12.5µM for 48h at 37°C and according to the manufacturer’s instruction form CellTiter-Glo® 2.0 Viability assay (Promega, G9242). Cytotoxicity was evaluated by measuring the activity of the lactate dehydrogenase (LDH) released in the supernatants of cells after treatment as seen above and according to the manufacturer’s instructions from the cytotoxicity detection kit (11644793001, Roche).

### 2.12. Bioenergetic analysis

Bioenergetic analyses were performed using the Seahorse XFe24 analyzer. Cells were seeded in a 24-well plate for Seahorse (100777-004, Agilent) 7 days before experiment, then cells were treated for 2 days with perhexiline at 10 μM. The day before the experiment, sensor cartridge from the Xfe24 Fluxpak (102340-100, Agilent) was immersed in the Seahorse XF calibrant solution (100840-000, Agilent) and incubated at 37°C without CO2 overnight. Seahorse medium (102353-100, Agilent) was prepared by adding pyruvate (1 mM, 103578-100, Agilent), L-glutamine (2 mM, 103579-100, Agilent) and glucose (10 mM, Agilent, 103577-100) and pH adjusted to 7.4. One hour before starting the analysis, cells were washed with the complete Seahorse medium and incubated for 45 min at 37°C in an incubator without CO2 while the analyzer was calibrating. Mitochondrial function analysis was performed using the Cell MitoStress Test kit (103015-100, Agilent). Measurements consisted in cycles of 3 min mix, 2 min wait and 3 min measure before and after oligomycin (2μM), carbonyl cyanide-4-phenylhydrazone (FCCP at 2μM) and antimycin A/rotenone (0.5μM) injections. Three measurement cycles of oxygen consumption rate (OCR) were performed at basal state, before any injection, and at metabolic stress state, simulated by the FCCP injection. Once the assay is completed, cells were fixed and stained with Hoechst for normalization.

### 2.13. β-oxydation of palmitic acid by mitochondria

Mitochondrial fatty acid oxidation was determined by measuring the acid-soluble radiolabeled metabolites resulting from mitochondrial oxidation of [U-^14^C] palmitic acid (PerkinElmer, Villebon-sur-Yvette, France). Briefly, adherent HepaRG cells in 96-well plate were incubated for 3 hours at 37°C in a red-free William’s E medium supplemented with 1 % fatty acid free BSA, 1 mM L-carnitine, [U-^14^C] palmitic acid (185 Bq/well), 100 μM cold PA and 1% DMSO. At the end of the incubation, 100 μL of 6 % perchloric acid was directly added in each well to promote precipitation of non-oxidized fatty acids. Then, cells were centrifuged at 2,000 g for 10 minutes at room temperature. After centrifugation, 130 μL of supernatant was collected and transferred in 4 mL of scintillation liquid. [U-^14^C] labeled acid soluble β-oxidation products were measured using a Tri-Carb 4910TR liquid scintillation counter (PerkinElmer, Villebon-sur-Yvette, France). Results were normalized to total protein content and expressed as counts per minute (CPM)/μg of proteins.

### 2.14. Statistical analyses

Results were expressed as mean ± SEM (the standard error of the mean) of at least three independent experiments. Comparisons between groups were performed using one-way ANOVA followed by a post hoc Dunnett’s test or Bonferroni’s test when the normality test was positive, and Mann-Whitney when data were not normally distributed. Graph Pad Prism 8 software (GraphPad Software, San Diego, CA, USA) was used for all statistical analyses) and the threshold for statistical significance was set to *p < 0.05, **p < 0.01, ***p < 0.001, ****p < 0.0001.

## III RESULTS

### 3.1. CYP2D6 expression and activity in parental and transgenic HepaRG cells

In our laboratories, the HepaRG cells are routinely expanded by successive passages every 2 weeks. When full differentiation is required, the cells are maintained 2 more weeks in appropriate medium to obtain metabolically competent hepatocyte-like HepaRG cells at days ∼30 after plating. Differentiated cells can also be frozen and stored in liquid nitrogen to provide ready to use functional hepatocyte-like cells after thawing (**Supporting information 1**).

In order to achieve efficient transduction with lentiviral particles encoding CYP2D6 and GFP, confluent 14 day-HepaRG cells were trypsinized and plated at low cell density and infected 24 h after seeding. Forty-eight h after infection of progenitor HepaRG cells, culture medium containing lentiviral particles was discarded and the transduced cell population were expanded for 2 weeks. Then, a working bank of frozen progenitor cells with stable expression of GFP was created (see Material and Method section). Both parental (wild-type, WT) and CYP2D6^+^ progenitor cells corresponding to HPR101 and HPR1LV01 references, respectively (**Supporting information 1**), were thawed and plated to monitor the cell morphology in phase contrast microscopy during proliferation and differentiation over 4 weeks (**Figure 1A**). In phase contrast microscopy, morphologies of CYP2D6^+^ versus parental progenitor cells appeared very similar at the different time points of the 30-day procedure of expansion and differentiation. In order to analyze the expression of CYP2D6 throughout the 30-day procedure of HepaRG cell differentiation, both parental and transduced cells were collected at different time points after plating and immunoblotting was performed to detect CYP2D6 as well as CYP3A4, which is a DME expressed at low levels in progenitor cells and strongly induced in differentiated HepaRG cells (**Figure 1B**). As expected CYP2D6 protein levels were too low to be detectable by immunoblotting in parental cells while a strong expression was found in HepaRG cells infected with lentiviral particles encoding CYP2D6 (**Figure 1B**). The CYP2D6 expression level did not appear significantly modified during the 30 days of culture using the semi-quantitative western-blot analysis.

**Figure 1:**
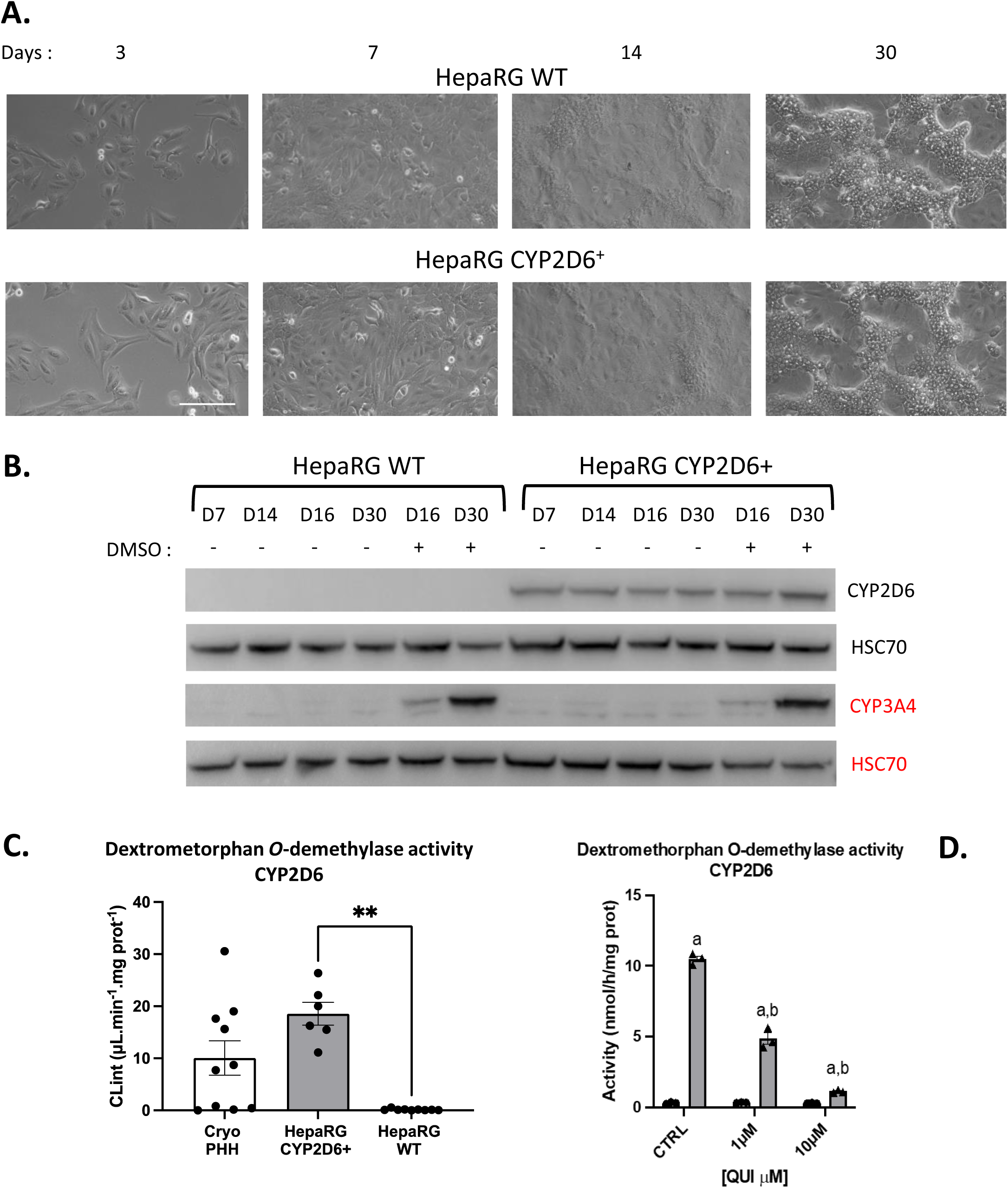
Expression and activity of CYP2D6 in the transgenic HepaRG cell line. A) Parental (WT) and CYP2D6^+^ progenitor cells were thawed and plated to monitor the cell morphology in phase contrast microscopy at days 3, 7 14 and 30 days after plating. B) Immunoblotting of CYP2D6 and CYP3A4 in parental (WT) and CYP2D6^+^ cells at different times after plating until days 30 in absence (-) or presence (+) of DMSO. HSC70 protein was used as loading control. C) Dextrometorphan *O*-demethylase activity (Cl_int_ values expressed in μL.min^-1^.mg^-1^) measured in human hepatocytes (PHH), HepaSH cells, parental (WT) and CYP2D6^+^ HepaRG cells. Statistics: *p < 0.05 and **p< 0.01 between Parental and CYP2D6^+^ HepaRG cells. D) Inhibition of dextrometorphan *O*-demethylase activity in CYP2D6^+^ HepaRG cells by quinidine at 1 and 10 μM. Statistics: *p < 0.05 between parental and CYP2D6^+^ HepaRG cells (a), and quinidine-treated cells versus untreated CYP2D6^+^ HepaRG cells (b).

Since the lentiviral particles used to transduced progenitor HepaRG cells also encoded GFP, we looked for the expression of this fluorescent marker during proliferation and differentiation of CYP2D6^+^ expressing cell population by fluorescence microscopy (**Supporting information 2A**) and flow cytometry (**Supporting information 2B**). In fluorescence microscopy, nearly all cells appeared to express GFP although the intensity of fluorescence varied between cells. In addition, the GFP fluorescence seemed to increase in a time-dependent manner over the 30 days of culture (**Supporting information 2A**). By flow cytometry, we demonstrated that more than 98% of cells were GFP positive (GFP^+^) at all days after plating with large differences in the overall means of fluorescence. These data indicated that the GFP expression levels varied among the HepaRG cell population and increased during the proliferation and/or differentiation process between days 3 and 30 after plating (**Supporting information 2B**). They also confirmed that transduced progenitor cells were able to differentiate in both hepatocyte- and cholangiocyte-like cells, which all expressed GFP.

We next investigated the CYP2D6 enzymatic activity in the whole differentiated HepaRG cell population including both hepatocyte- and cholangiocyte-like cells (**Figure 1C**). We found a high Cl_int_ values (18 μL.min^-1^.mg^-<u>1</u>^) in CYP2D6^+^ HepaRG cells compared to parental wildtype cells for which the activity was very low (0.1467 μL.min^-1^.mg^-1^). These values were compared with those obtained for human hepatocytes obtained by dissociation of liver biopsies, cryopreserved at Biopredic^International^, thawed and used in suspension for CYP2D6 activities. The mean Cl_int_ values obtained with cryopreserved human hepatocytes was 10.05 μL.min^-1^.mg^-1^ coupled with a noticeably high variability between donors (range from 0 to 30 μL.min^-1^.mg^-1^) (**Figure 1C**), demonstrating that CYP2D6^+^ HepaRG cells expressed a CYP2D6 catalytic activity in the range of those found in PHH.

To verify whether the catalytic activities measured CYP2D6^+^ HepaRG cells were indeed due to the enforced expression of the CYP2D6, we used quinidine, a potent competitive inhibitor of CYP2D6 (**Figure 1D**). As expected, CYP2D6 catalytic activities were inhibited in a dose-dependent manner by quinidine.

Since we had shown that GFP was expressed in both hepatocyte- and cholangiocyte-like HepaRG cells (**Supporting information 2A, B**), we next studied the expression levels of both GFP and CYP2D6 in these two cell populations when fully differentiated at 30 days after plating (**Figure 2**). To address this question, we took advantage of the procedure established in our laboratory to selectively detach both hepatocyte- and cholangiocyte-like HepaRG cells by mild trypsin incubation (**Figure 2A**). In microscopy, the GFP fluorescence in hepatocyte-like cells first appeared more intense compared to that observed in cholangiocytes (**Figure 2A**) but the quantification by flow cytometry confirmed that both cell populations were GFP positive (∼100%) and demonstrated that means of fluorescence were not statistically different between hepatocyte- and cholangiocyte-like HepaRG cells (**Figure 2B**).

**Figure 2:**
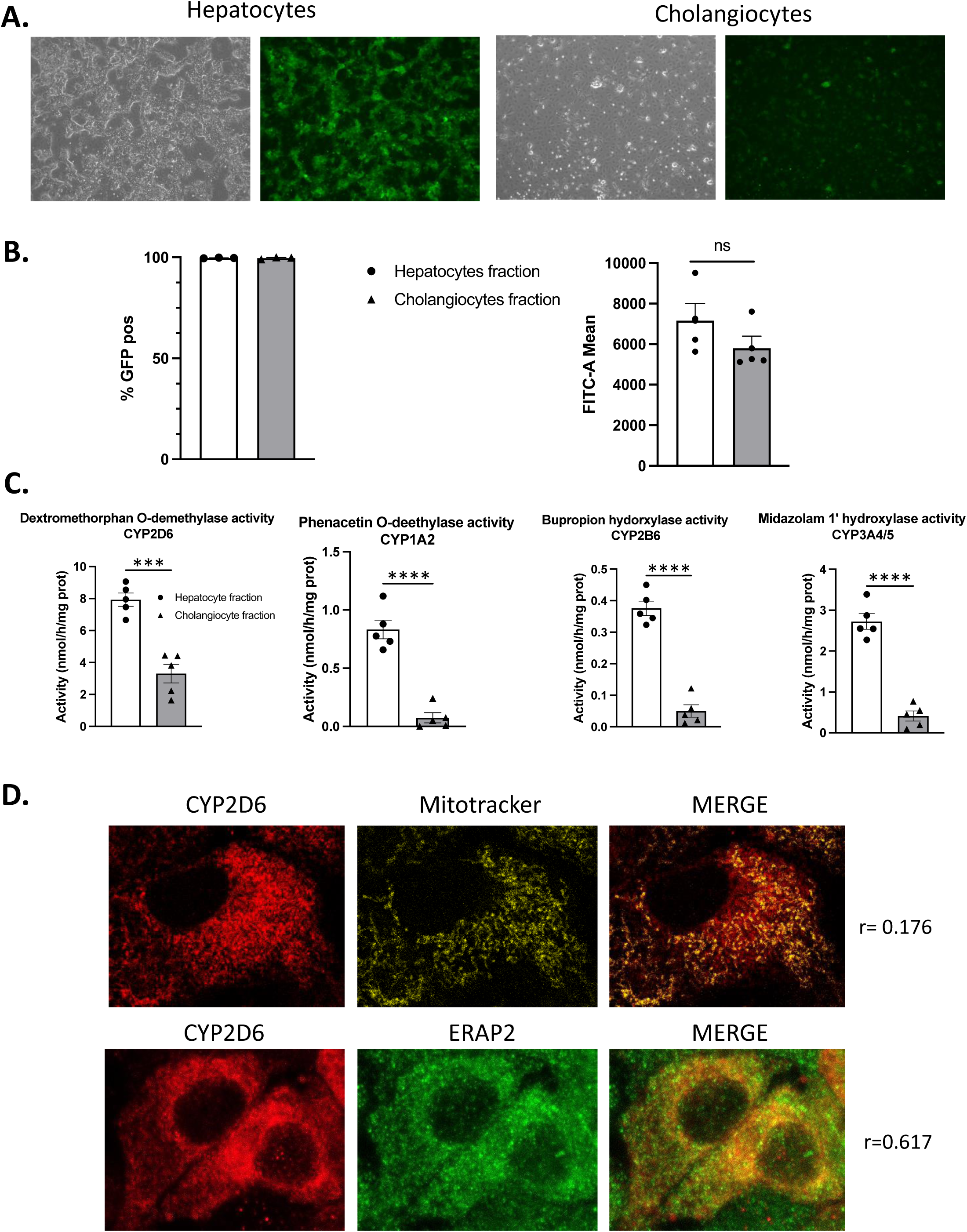
Expression of GFP and CYP2D6 in differentiated hepatocyte- and cholangiocyte-like HepaRG cells. *In situ* detection of GFP by fluorescence microscopy (A) and by flow cytometry (B) in hepatocyte- and cholangiocyte-enriched cell populations. For flow cytometry data (B), results are expressed in percentages of GFP positive (Pos) cells and means of fluorescence (FITC-A mean) in both hepatocyte- and cholangiocyte-enriched cell populations. C) Dextrometorphan *O*-demethylase, phenacetin O-deethylase, bupropion hydroxylase and midazolam 1’-hydroxylase activities probing CYP2D6, CYP1A2, CYP2B66 and CYP3A4/5, respectively, in both hepatocyte- and cholangiocyte-enriched cell populations. All data were significantly different between the two cell populations: ***p < 0.001, ****p < 0.0001. D) *In situ* detection of CYP2D6 (red staining) and ERAP2 (green staining) by immunofluorescence and mitochondria staining using mitotracker.

In these two cell populations, we then measured CYP1A2, CYP2B6, CYP3A4/5 and CYP2D6 enzymatic activities (**Figure 2C**). CYP1A2, CYP2B6, CYP3A4/5 catalytic activities were much higher in hepatocyte-like cell cultures compared to those measured in cholangiocyte fraction (∼8-fold difference), which confirmed that we had obtained enriched populations for both cell types. For dextromethorphan O-demethylase CYP2D6 activity, we found that hepatocyte-like cells had ∼2.5-fold higher activities than the values obtained in cholangiocytes (**Figure 2C**). These data demonstrated that both hepatocyte- and cholangiocyte-like cells expressed CYP2D6 but that the enzymatic activity was significantly higher in hepatocytes compared to cholangiocytes.

We next studied the intracellular localization of the CYP2D6 in transduced hepatocyte-like cells by confocal immunofluorescence (**Figure 2D**). In this experiment, we also used the gree fluorescent mitotracker and the detection of the Endoplasmic Reticulum Aminopeptidase 2 (ERAP2) staining the mitochondria and endoplasmic reticulum (ER), respectively, to determine the subcellular localization of the CYP2D6. As expected, a strong CYP2D6 staining was observed in transgenic HepaRG cells, which poorly co-localized with the fluorescent mitotracker while the co-staining with ERAP2 was relatively high (**Figure 2D**) suggesting that CYP2D6 was mainly localized in the ER.

### 3.2. Tramadol biotransformation and perhexiline toxicity in CYP2D6 transgenic HepaRG cells

Since we had established a stable CYP2D6 expressing HepaRG cell line, we next compared the metabolism of tramadol in both parental and CYP2D6^+^ progenitor and differentiated HepaRG cells (**Figure 3**). CYP2D6, CYP3A4 and CYP2B6 catalyze the metabolism of tramadol (Miotto et al., 2017) with the productions of *O*-desmethyltramadol (metabolite M1) via the *O*-demethylation reaction catalyzed by CYP2D6 and *N*-desmethyltramadol (M2) catalyzed by CYP2B6 and CYP3A4 (**Figure 3A**). Then, from these two M1 and M2 metabolites, *N*,*O*-didesmethyltramadol (M5) is produced by CYP2B6/CYP3A4 and CYP2D6, respectively. Alternatively, *N*,*N*-bisdesmethyltramadol (M3) can be generated from M2 by CYP2B6 and CYP3A4 further converted into *O*-desmethyl-*N*,*N*-bisdesmethyltramadol (M4).

**Figure 3:**
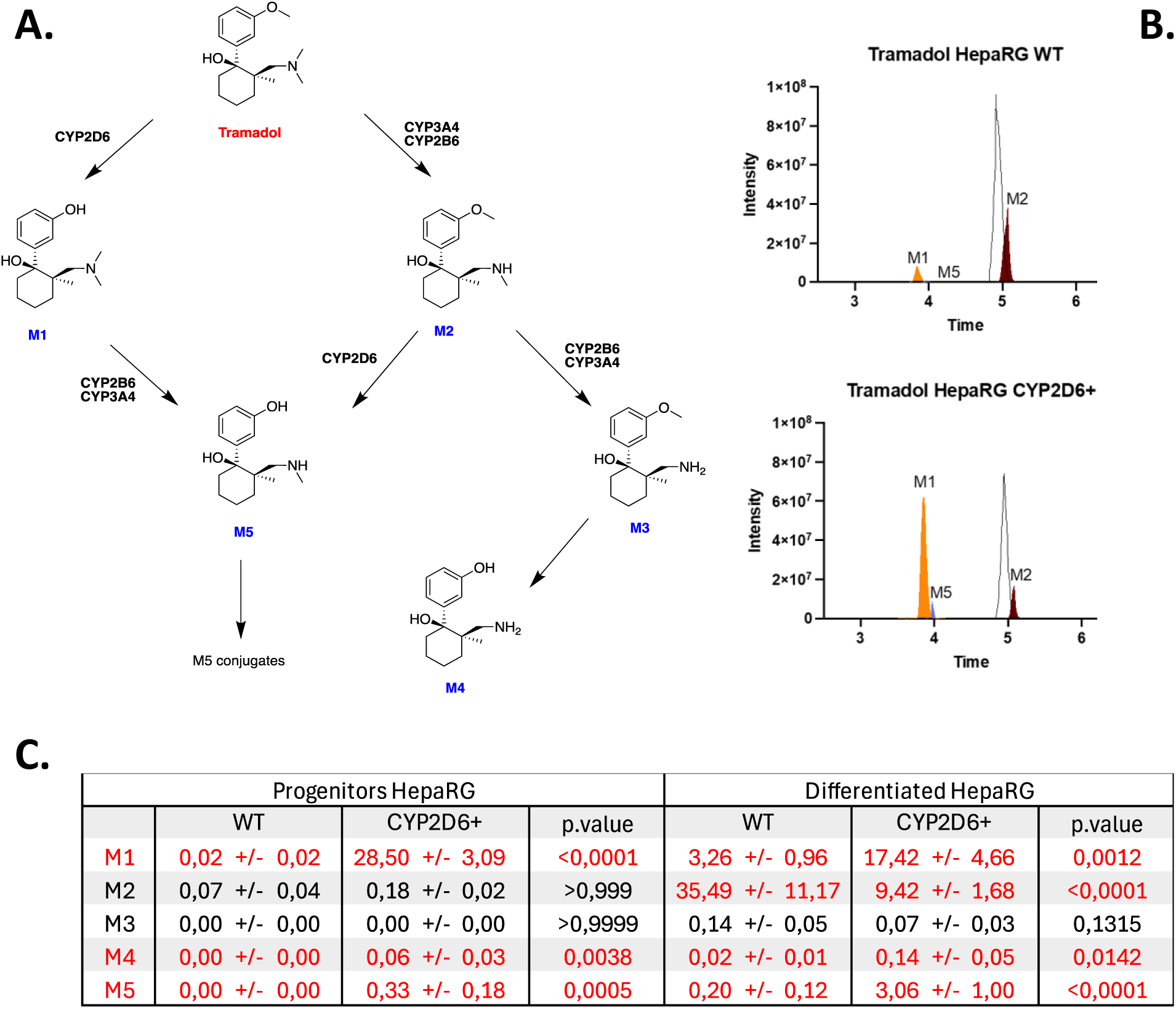
Tramadol biotransformation in parental and CYP2D6^+^ HepaRG cells. A) Schematic representation of the biotransformation of tramadol via CYP2D6, CYP2B6 and CYP3A4 enzymatic activities and the main metabolites of tramadol: M1: *O*-desmethyltramadol, M2: *N*-desmethyltramadol, M3: *N*,*N*-bidesmethyltramadol, M4: *O*-desmethyl-*N*,*N*-bisdesmethyltramadol, M5: *N*,*O*-didesmethyltramadol (M1, M2, M3, M4 and M5). B) Chromatograms of M1-M2 metabolites detected in parental (HepaRG WT) and CYP2D6^+^ transgenic HepaRG cells. C) Table presenting the quantification LC–MS/MS analysis of metabolites M1 to M5 in progenitor and differentiated parental (WT) and CYP2D6^+^ HepaRG cells. Results were expressed in µg/L.

In serum of humans, both M1 and M2 metabolites can be quantified by LC-MS/MS (**Supporting Information 6**) with various M1/M2 ratios most likely correlating with the CYP2D6, CYP3A4 and CYP2B6 expression levels and catalytic activities. In culture media of parental differentiated HepaRG cells incubated with 20µM tramadol, we found high concentrations of M2 (35.49±11.17 µg/L) and low amounts of M1 (3.26±0.96 µg/L) (**Supporting Information 3, Figure 3B, C**). When using progenitor CYP2D6^+^ HepaRG cells, only M1 metabolite was detected in culture media while M1 (17.42±4.66 µg/L), M2 (9.42±1.68 µg/L) and M5 (3.06±1 µg/L) metabolites were found in media of differentiated cells (**Figure 3B, C**).

These results demonstrated that expression of CYP2D6 in differentiated HepaRG cells allowed the production of the main metabolites of tramadol in concentrations detectable by LC-MS/MS in culture media.

We also studied whether the expression of CYP2D6 in HepaRG cells could modify significantly the cytotoxicity of the perhexiline (**Figure 4**), a prophylactic antianginal agent known to induce severe adverse effects including hepatotoxicity involved mitochondrial damages and apoptosis (Ren et al., 2022). The major route of perhexiline metabolism in humans is hydroxylation (Sørensen et al., 2003) mainly by CYP2D6 producing two main metabolites, the cis and trans isomers of hydroxyperhexiline, although CYP1A2, CYP2C19, and CYP3A4 also contribute to the metabolism of perhexiline (Ren et al., 2022). It has been previously shown that the cytotoxicity of perhexiline was reduced significantly in CYP2D6-overexpressing HepG2 cells, in comparison to the control HepG2 cells while enforced expression of CYP1A2, CYP2C19, and CYP3A4 had no effect on the toxicity of this drug (Ren et al., 2022). Parental and CYP2D6^+^ differentiated HepaRG cells were exposed to various concentration of perhexiline and the cell viability (relative ATP contents) and cytotoxicity (LDH release) were measured (**Figure 4A**). We found that relative ATP contents were lower and LDH release was higher in parental HepaRG cells in comparison to those measured in CYP2D6^+^ cells. Consequently, the IC_50_ was higher in CYP2D6^+^ versus parental cells (**Figure 4B**). Since the toxicity of perhexiline involves mitochondrial damages, we studied mitochondrial parameters in HepaRG cells exposed to this drug and found that ATP production and mitochondrial coupling were higher while proton leak was lower in CYP2D6^+^ cells compared to parental cells (**Figure 4C**). In addition, the levels of mitochondrial β-oxidation of linoleic acid was also lower in parental cells compared to that in CYP2D6^+^ cells expose to perhexiline at 20μM (**Figure 4D**). These data demonstrated that the enforced expression of CYP2D6 metabolizing perhexiline partially protected HepaRG cells from the toxicity of this drug for concentration below the IC_50_.

**Figure 4:**
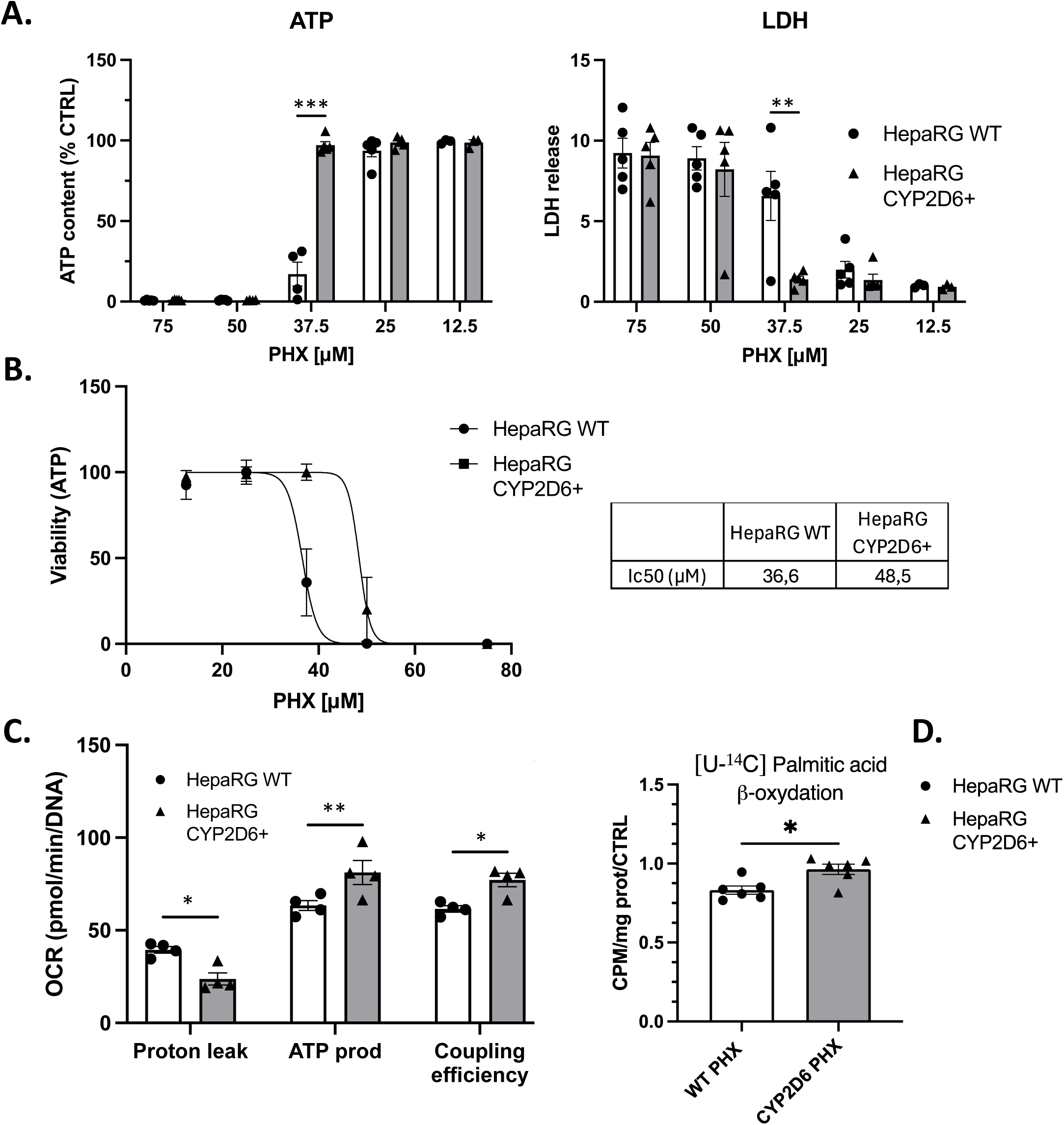
Perhexiline toxicity in parental and CYP2D6^+^ HepaRG cells. A) Relative ATP contents (left panel) and LDH release (right panel) and B) IC_50_ in parental (WT) and CYP2D6^+^ differentiated HepaRG cells exposed to various concentration of perhexiline. C) Oxygen Consumption Rate (OCR) in parental (WT) and CYP2D6^+^ differentiated HepaRG cells exposed to perhexiline at 10μM and quantifications of proton leak, ATP production and coupling efficiency (left panel) and D) levels of mitochondrial β-oxidation of linoleic acid. All data were significantly different between the two cell lines: *p < 0.05 and **p< 0.01

### 3.3. Impact of the lentiviral insertion and CYP2D6 expression on proliferation and differentiation of transgenic HepaRG cells

The monitoring of parental and CYP2D6 HepaRG cell morphologies during 30 days after plating suggested that proliferation and differentiation were not altered (**Supporting information 2A**). To evaluate more deeply the impact the lentiviral transgene integration and CYP2D6 expression on the recombinant HepaRG cell fate, we performed a series of experiments characterizing their proliferation and differentiation.

First, we compared the proliferation rates between parental (GFP negative, GFP^-^) and CYP2D6^+^/GFP^+^ cells (**Supporting information 4A**). To address this issue, we mixed both GFP^-^ parental and CYP2D6^+^/GFP^+^ cells in various proportions (30%/70%, 50%/50% and 70%/30%) and we monitored the percentages of GFP^-^ versus GFP^+^ cells at different times points after plating (**Supporting information 4A**) and over several passages (data not shown). The rationale of this experiment was the following: If one of the 2 cell populations grew faster, the percentages of the faster-growing cell population would increase during expansion. By flow cytometry, we showed that the percentages of both GFP^-^ and GFP^+^ cells were not altered during the proliferation period during the first 14 days after plating (**Supporting information 4A**) demonstrating that CYP2D6^+^/GFP^+^ cells had a proliferation rate similar to their parental counterparts.

Then, we performed gene profiling by Bulk RNA barcoding and sequencing (BRB-seq) technology using RNAs prepared from both parental and CYP2D6^+^/GFP^+^ cells (**Figure 5**) at different steps of the proliferation and differentiation process and in cryopreserved differentiated cells that were cultured for 2 weeks after thawing. We selected the following times points: days 2 and 4 after plating when progenitor cells are actively proliferating, day 7 when proliferation reduces and cells begin committing to hepatocyte or cholangiocyte lineages, at day 14 when cells are quiescent and committed to the two lineages, at day 32 after cell thawing of differentiated cells and at days 37 and 44, when cells are fully differentiated (**Supporting information 1**).

**Figure 5:**
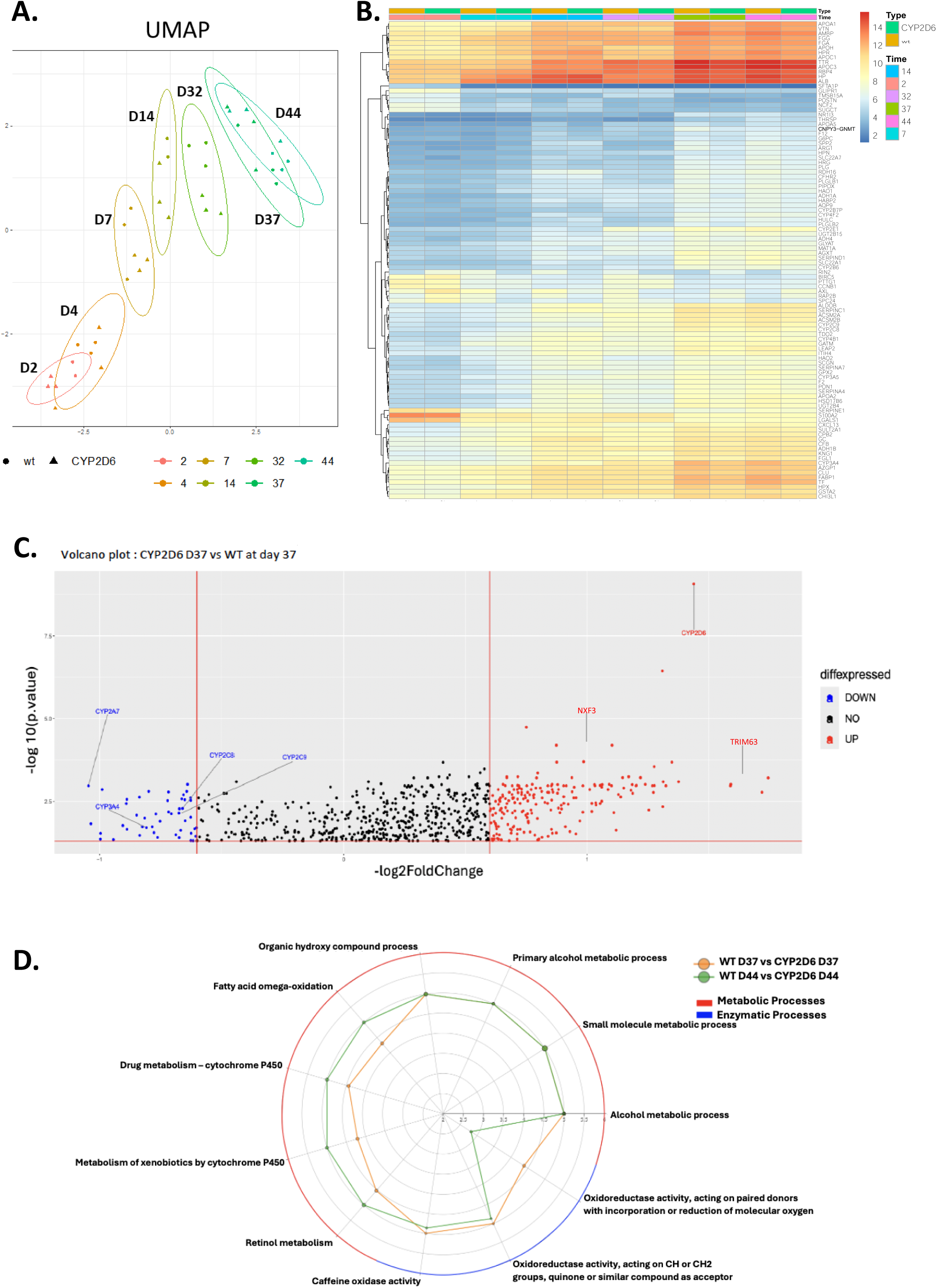
Comparative BRB-seq analysis of parental (wild-type, WT) and CYP2D6^+^ HepaRG Cells during proliferation and differentiation. A) UMAP representation of parental (WT) and CYP2D6^+^ HepaRG cells at different time points after plating (2, 4, 7, 14, 32, 37 and 44 days). The first dimension is mainly associated to the time of culture. B) Heatmap of the top 100 most differentially expressed genes during differentiation, with consistent expression patterns observed between parental (WT and CYP2D6^+^ HepaRG cells. C) Volcano plot of genes differentially expressed at D37 highlighting significant downregulation of various CYP450 genes in transgenic cells compared to parental cells. D) Radar plot showing enrichment of gene ontology (GO) terms related to xenobiotic metabolism at D37 and D44. The size of the points is the ratio r/R and the scale corresponds to the log10 of the adjusted p-value, where r = number of genes in the list associated with the term of interest and R = total number of genes associated with one term of interest.

The genes that were found differentially expressed between parental and CYP2D6^+^/GFP^+^ cells at each time point were analyzed using Uniform Manifold Approximation and Projection (UMAP) to draw a two-dimension representation of the different data sets in order to determine whether the parental and CYP2D6^+^/GFP^+^ cells belonged to the same cluster during the differentiation process (**Figure 5A**). We found that the two cell lines segregated in the same clusters during proliferation and differentiation since the first dimension is mainly associated to the time of culture after plating indicating that their overall gene expressions were very similar in the two lines. To further confirm this statement, the top 100 genes showing the highest differential expression have also been visualized in a heatmap (**Figure 5B**). Similarly, the heatmap evidenced that gene expression in parental and CYP2D6^+^/GFP^+^ cells clustered at each time point. When considering the genes that showed the highest increase in expression levels as a function of time after cell plating, numerous genes encoding hepatocyte specific functions were found including albumin, haptoglobin, phase I and II enzymes (**Figure 5B**). In contrast, we also found some genes that showed a strong decrease in their expression levels such as genes identified in hepatocellular carcinomas including SFTA1P encoding a long non-coding RNA (Liao et al., 2018), the extracellular matrix periostin (Chen et al., 2017) and NCF2 (Huang et al., 2023), and CCNB1 encoding the cell cycle regulator cyclin B1 (**Figure 5B**).

Considering that the overall gene profiling between parental and CYP2D6^+^ cells demonstrated that transgenic HepaRG cells had kept the ability to differentiate into hepatocyte- and cholangiocyte-like cells, we compared the induction of CYP1A2, CYP2B6 and CYP3A4/5 catalytic activities in both parental and CYP2D6^+^ differentiated cells treated with canonical inducers omeprazole (CYP1A2), rifampicin (CYP3A4) and phenobarbital (CYP2B6) (**Supporting information 4B**). We found that the induction of CYP1A2 and CYP3A4/5 catalytic activities were similar in the two cell lines while CYP2B6 enzymatic activity was higher in CYP2D6^+^ cells compared to the parental cells. Similarly, we studied parameters of mitochondrial respiration using the Seahorse technology (**Supporting information 4C**) since the differentiated HepaRG cells have been shown to exhibit a O_2_ consumption, high oxidative phosphorylation and ATP production (Peyta et al., 2016 ; Daniel et al., 2024). The O_2_ consumption rate, basal and maximal respiration, proton leak and ATP production were nearly identical between parental and transgenic HepaRG cells.

We next looked for genes that were differentially expressed between parental and CYP2D6^+^ cells at days 37 and 44 when cells are fully differentiated (**Supporting information 1**). Using Volcano plots (**Figure 5C**), we identified a list of genes that were either up or down-regulated in differentiated CYP2D6^+^ versus parental cells (**Supporting Information 4**). At day 37, 195 genes were up-regulated while 44 genes were down-regulated in CYP2D6^+^ versus parental cells. At day 44, 250 genes were up-regulated and 61 genes were down-regulated. As expected, CYP2D6 was found up-regulated in the transgenic cell line at 37 and 44 days. Interestingly, when comparing the top ∼20 genes showing the highest differences in fold change only three down-regulated and 8 up-regulated genes were found in common at 37 and 44 days (**Supporting Information-Table 4**). Radar plot, however, showed enrichment of gene ontology (GO) terms related to lipid, xenobiotic and drug metabolisms and oxidoreductase activities at days 37 and 44, (**Figure 5D**). Among xenobiotic and drug metabolism genes, expressions of several phase I enzymes were significantly decreased, especially CYP2A7, CYP2C8/9 and CYP3A4 (**Supporting Information Table 4**). This unexpected data led us to hypothesize that either the insertion of the lentiviral transgene and/or the high CYP2D6 expression levels may have contributed to significantly affect the expression of these phase I proteins.

To address this question, we knocked-down the expression of the CYP2D6 within the lentiviral transgene using the CRISPR/Cas9 technology (**Figure 6A, Supporting Information 5**) and we studied the mRNA expression levels of these CYP by RT-qPCR in parental and transgenic cell lines (**Figure 6B**). The invalidation of the CYP2D6 encoding sequence within the lentiviral transgene also knocked-down the GFP expression, which was used to sort the GFP negative cells by FACS and to derive a KO^CYP2D6/GFP^ HepaRG cell line with ∼4% of residual GFP^+^ cells (**Figure 6A**). By RT-qPCR, we showed that the CYP2D6 mRNA levels were strongly reduced in the KO^CYP2D6/GFP^ HepaRG cell line confirming the invalidation of the CYP2D6 lentiviral transgene. In contrast, we did not show significant differences in the mRNA levels of CYP2A7, CYP2C8/9 and CYP3A4 (**Figure 6B**). We also studied the mRNA of transcription factors involved in the regulation of the CYP gene expression. Unexpectedly, we found that AhR, FXR, PXR, HNF4a and SHP mRNA levels were also decreased in the CYP2D6/GFP transgenic cells compared to those levels in the parental cells and that their expression was higher in the KO^CYP2D6/GFP^ HepaRG cell line with mRNA levels similar to those measured in the parental cells.

**Figure 6:**
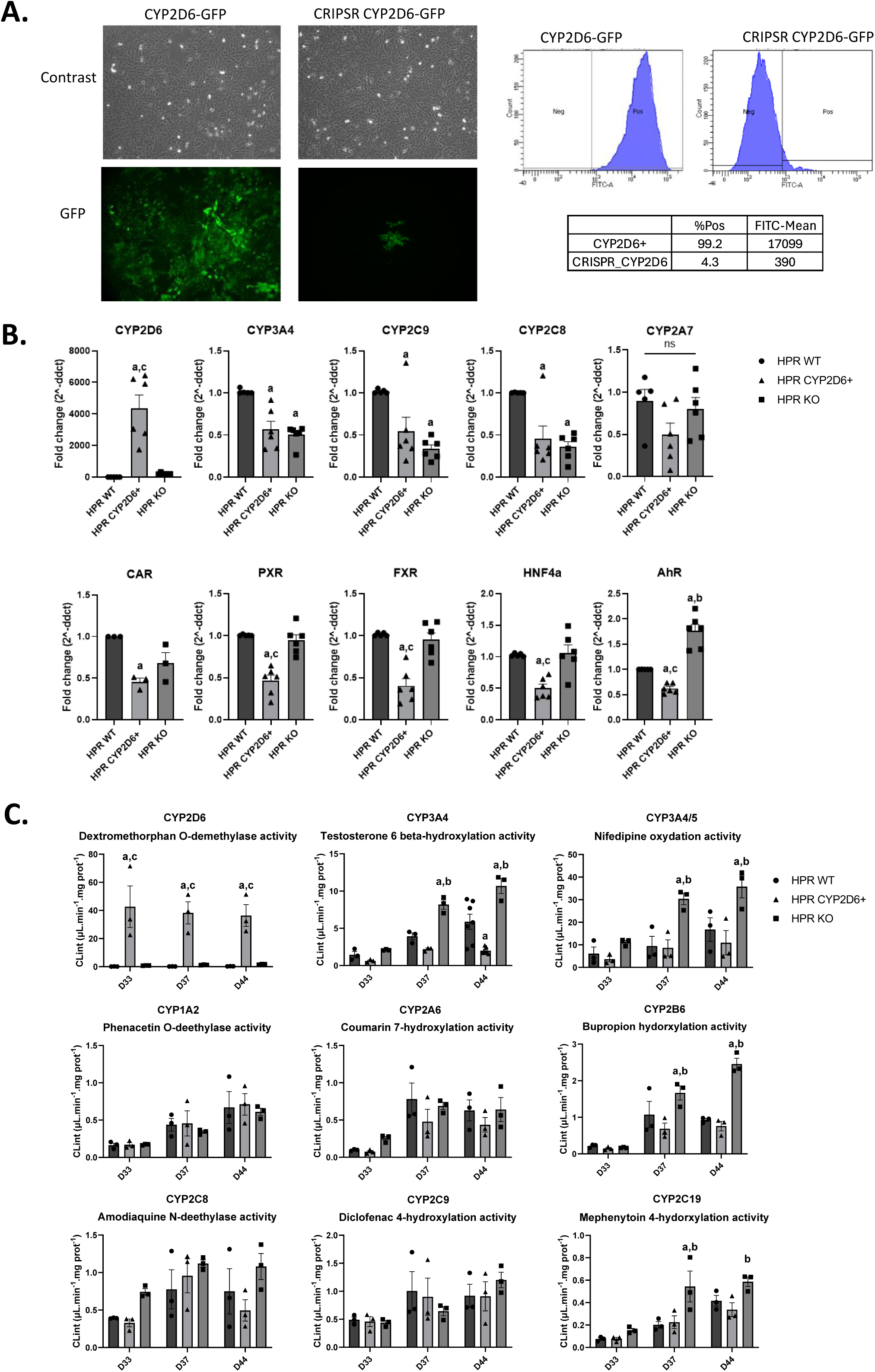
Expression and activity of various CYP450 in parental, CYP2D6^+^ and KO^CYP2D6/GFP^ HepaRG cells. A) Detection of GFP by fluorescence microscopy and by flow cytometry in CYP2D6^+^ and KO^CYP2D6/GFP^ HepaRG cells. Results are expressed in percentages of GFP positive (Pos) or negative (Neg) cells and means of fluorescence (FITC-A mean). B) RT-qPCR evaluating the relative mRNA expression of CYP2D6, 3A4, 2C8, 2C9, 2A7, AhR, FXR, PXR, HNF4α and SHP in parental (HRP WT), CYP2D6^+^ and KO^CYP2D6/GFP^ (HRP KO) HepaRG cells. Statistics: *p < 0.05 between parental (HRP WT = a), CYP2D6^+^ (b) and KO^CYP2D6/GFP^ (HRP KO = c) HepaRG cells.

We next measured several phase I catalytic activities in the three HepaRG cell lines at 33, 37 and 44 days of culture corresponding to two weeks of culture after thawing (**Figure 6C**). As expected CYP2D6-mediated dextromethorphan O-demethylase activity was very high in GFP^+^ cells and very low in parental and KO^CYP2D6/GFP^ HepaRG cell lines, confirming that the CRISPR/Cas9 procedure invalidated the CYP2D6 activity in most cells. The other enzymatic activities that we quantified increased between days 33 and 44 (**Figure 6C**) concomitantly with the acquisition of the highest level of differentiation (**Figure 5A, B**). Interestingly, we found that the testosterone 6β-hydroxylation catalyzed by CYP3A4 was significantly reduced in CYP2D6^+^ HepaRG cells compared to that measured in parental cells while the same activity was higher in KO^CYP2D6/GFP^ HepaRG cells (**Figure 6C**). Similarly, we found that bupropion hydroxylase CYP2B6-, mephenytoin 4-hydroxylase CYP2C19- and nifedipine oxidation CYP3A4/5-dependent activities were higher in KO^CYP2D6/GFP^ cells compared to those found in parental and CYP2D6^+^ HepaRG cells (**Figure 6C**). In contrast, CYP2C8, CYP2C9, CYP1A2 and CYP2A6 dependent activities were not significantly different between the three cell lines. Together, these data showed that the process of lentiviral transduction and/or enforced expression of CYP2D6 and GFP in HepaRG cells modulated the mRNA levels of several CYP450 genes as well as nuclear receptors regulating these CYP450. Invalidation of the CYP2D6 and GFP expression rescued the mRNA levels of these nuclear receptors but not the CYP450 mRNA levels. In addition, we did not observe a correlation between these CYP450 mRNA levels and the corresponding CYP450 catalytic activities.

Interestingly, the transcriptomic experiments showed also highly up-regulated genes, which are not reported to be expressed in the normal liver but known to be regulated by viral infections (), including NXF3 and TRIM63 (**Figure 5C**). We then postulated that the WPRE inserted in the lentiviral construct (**Figure 7A**), which encodes a post-transcriptional regulatory element from the woodchuck hepatitis virus present in the CYP2D6-GFP encoding transcript, could generated the induction of NXF3 and TRIM63 rather than the indirect insertion of the lentiviral transgene into the cell genome. We next performed RT-qPCR analysis of these genes and we found a strong induction of WPRE, TRIM63 and NXF3 in CYP2D6^+^ HepaRG cells at D37 or D44 after lentiviral induction (**Figure 7B**). In addition, in KO^CYP2D6/GFP^ cells, the expression of WPRE, TRIM63 and NXF3 was silenced as observed for both GFP and CYP2D6, confirming the correlation between the expression of TRIM63 and NXF3 genes and the expression of the single transcript encoding GFP and CYP2D6, which also contains the non-coding WPRE sequence. These data strongly suggest that TRIM63 and NXF3 are induced by WPRE in differentiated HepaRG cells model and could participate to the modulation of gene and protein expression of hepatic functions.

**Figure 7:**
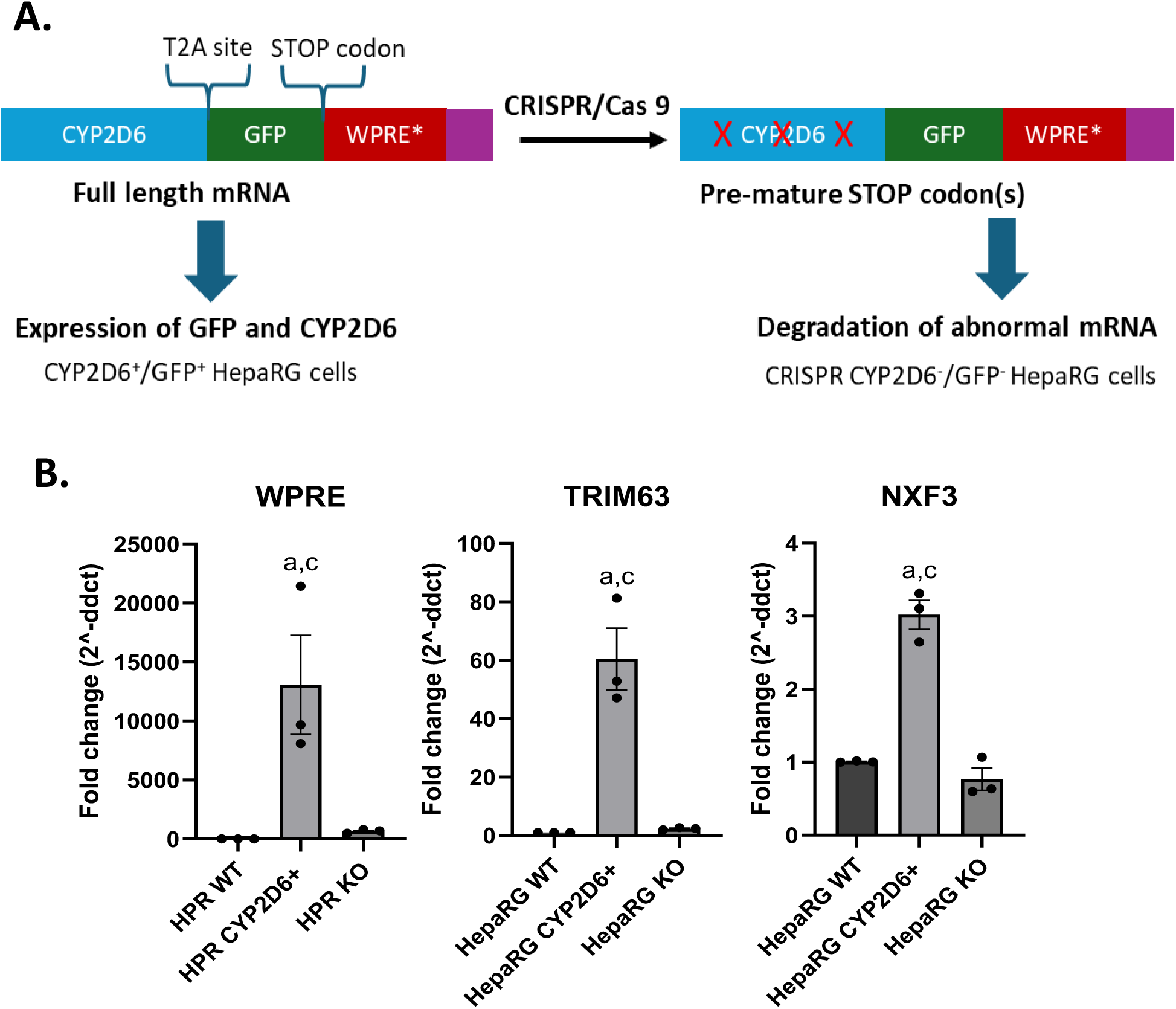
GFP and CYP2D6 lentiviral derived, NXF3 and TRIM63 mRNA expressions in parental CYP2D6^+^ and KO^CYP2D6/GFP^ HepaRG cells. A) Schematic representation of the lentiviral derived mRNA transcribed from the transgene inserted into the genome of HepaRG cells and CRISPR/Cas9 mediated alteration of the lentiviral transgene in KO^CYP2D6/GFP^ (HRP KO) HepaRG cells. B) RT-qPCR evaluating the relative mRNA expression of the CYP2D6-GFP-WPRE lentiviral derived transcript and NXF3 and TRIM63 in CYP2D6^+^ and KO^CYP2D6/GFP^ HepaRG cells. Statistics: *p < 0.05 between parental (HRP WT = a), CYP2D6^+^ (b) and KO^CYP2D6/GFP^ (HRP KO = c) HepaRG cells.

## IV DISCUSSION

Herein, we report the establishment of a transgenic HepaRG cell line expressing high levels of CYP2D6 using lentiviral transduction of progenitor cells without cell cloning or antibiotic selection to preserve the ability of HepaRG cells to proliferate and to differentiate into both hepatocytes and biliary cells. These transgenic cells exhibit a high CYP2D6 catalytic activity in the same range of those found in high metabolizers using cryopreserved human hepatocytes and HepaSH cell model, which are human hepatocytes expanded in an immuno-incompetent mice strain (Hasegawa et al., 2011 ; Zerdoug et al., 2024).

These data confirm that enforced expression of genes in HepaRG cells via lentiviral transduction, at least for genes that do not compromise either proliferation or survival, is very efficient when the infection is performed using progenitor cells. We previously used the same strategy to generate a recombinant HepaRG cell line expressing higher levels of CYP2E1 compared to that detected in the parental cells and a catalytic activity similar to what is measured in PHH (Quesnot et al., 2018). This enhanced CYP2E1 enzymatic activity allowed us to identify new metabolites of a specific CYP2E1 substrate, the chlorzoxazone, used to measure its activity *in vitro* and *in vivo*. These two transgenic HepaRG cells provide genetic and metabolic diversities from the parental HepaRG that can be used to compare biotransformation and cytotoxicity of xenobiotics between the initial HepaRG cells and transgenic counterparts. In this context, we demonstrated that the enforced expression of CYP2D6 in HepaRG cells better recapitulated the metabolism of tramadol found *in vivo* with the production of its two main metabolites, the *O*-desmethyltramadol and *N*-desmethyltramadol, in transgenic cells while parental produced *N*-desmethyltramadol only via the CYP2B6 and CYP3A4 activities. Similarly, we found that the cytotoxicity of perhexiline was reduced in CYP2D6 expressing cells compared to that observed for parental cells.

We next characterized in greater details the proliferation and differentiation of the transgenic CYP2D6 expressing HepaRG cell line. The ability of CYP2D6^+^/GFP^+^ HepaRG cell line to proliferate with a doubling time that appeared similar to parental cells was achieved by mixing the CYP2D6^+^/GFP^+^ and parental cells and demonstrating that the ratio between the two population did not change over time. In addition, we created a working bank of CYP2D6^+^/GFP^+^ cells by expansion of these cells through several passages further supporting that transgenic cells have kept the proliferation status of parental cells. Regarding the differentiation, we have gathered several evidences that the transgenic cells had also kept their ability to progress from hepatic progenitor cells towards well-differentiated hepatocyte-like cells. Data of BRBseq indeed indicated that both parental and CYP2D6^+^/GFP^+^ HepaRG cells sequentially expressed similar sets of genes characterizing the differentiation process described previously in numerous articles (Dubois-Pot-Schneider et al., 2014 ; Fekir et al., 2019 ; Dubois-Pot-Schneider et al., 2022). These data obtained at the mRNA level were confirmed with protein expressions and enzymatic activities, which demonstrated that major phase I activities were found at levels comparable to those measured in parental cells. Altogether, our results demonstrated that CYP2D6 expressing HepaRG cells proliferate and differentiate in a very similar manner that parental cells and that the lentiviral transgene insertion in HepaRG cell genome did not compromise these processes as we previously observed for other lentiviral infections in HepaRG cells (van der Mark et al., 2017 ; Quesnot et al., 2018 ; Pivert et al., 2019 ; Vlach et al., 2019 ; Heintze et al., 2021) and primary cultures of human hepatocytes (Zamule et al., 2008).

Nevertheless, we also identified from the data of BRBseq a panel of genes that were either up- or down-regulated in CYP2D6^+^/GFP^+^ HepaRG cells compared to gene expression in parental cells. We particularly focused on genes that were differentially expressed at days 37 and 44, the time-points corresponding to differentiated cells. Among the down-regulated genes, CYP2A7 and CYP3A4 expression levels were significantly reduced. This data led us to further evaluate the impact the lentiviral transgene integration and CYP2D6^+^/GFP^+^ expression on a panel of CYP450 both at the mRNA levels and enzymatic activities. In addition to parental and CYP2D6^+^/GFP^+^ cells, we generated KO^CYP2D6/GFP^ HepaRG cells, which kept the lentiviral insertion into their genome but lacked both CYP2D6 and GFP expressions. We confirmed that CYP3A4, CYP2C9, CYP2C8 and CYP2A7 mRNA levels were decreased in CYP2D6^+^/GFP^+^ cells compared to parental cells and knocking-down expression of CYP2D6 and GFP in KO^CYP2D6/GFP^ cells did not rescue this decrease. Unexpectedly, while the mRNA levels of transcription factors AhR, FXR, PXR, HNFA4 and SHP regulating CYP expression were also decreased in CYP2D6^+^/GFP^+^ cells, their expressions were higher in KO^CYP2D6/GFP^ cells. In addition, enzymatic activities of CYP2B6, 2C8, 2C9, 2C19, 1A2 and 2A6 were not significantly different between parental and CYP2D6^+^/GFP^+^ cells indicating that mRNA levels were not strictly correlated to enzymatic activities. In contrast, we confirmed that CYP3A4 enzymatic activities were lower in CYP2D6^+^/GFP^+^ cells compared to parental cells and that the invalidation of CYP2D6^+^/GFP in KO^CYP2D6/GFP^ HepaRG cells totally rescued the CYP3A4 activities. These last data suggested that the high expression levels of CYP2D6 and GFP would negatively affect the CYP3A4 activity. This observation may partially be explained by data indicating that different CYP450 subunits interact with each other to form heteromeric complexes, which functionally impact individual CYP450 activities (Davydov, 2011; Reed and Backes, 2016). This functional interaction has been suggested in CYP3A4/CYP2D6 double transgenic humanized mice (Felmlee et al., 2008) and, more recently, it has been reported that enzymatic activity of CYP3A4 was dependent on the composition in all CYP450 (Dangi et al., 2021).

The rationale for the comparison of gene expressions and enzymatic activities in parental, CYP2D6^+^/GFP^+^ and KO^CYP2D6/GFP^ cells was to attempt to discriminate between the impact of lentiviral insertion into the host cell genome and enforced expression of CYP2D6. In this context, there is an abundant literature describing the mechanisms of lentiviral insertion in human cell genome and the genotoxicity resulting from this insertion (Gil-Farina and Schmidt, 2016 ; Biasco et al., 2017). The lentiviral integration is influenced by the chromatin landscape encountered during nuclear entry of the viral genome and involves cellular proteins including transcription factors, which bind to virus long terminal repeats (LTR) and virus pre-integration complexes (PICs) to select the insertion site(s) (Suleman et al., 2022) preferentially in transcriptional active regions (Craigie and Bushman, 2014). Thus, integration can occur in many different sites depending on the status of differentiation and proliferation of the host genome. The integration may lead to insertional mutagenesis either disrupting normal expression of one or several genes in one cell and/or regulatory mechanisms resulting in aberrant differentiation pattern, malignant transformation or even apoptosis. In our experimental setting, we used a multiplicity of infection close to 1 in order to reduce the number of cells with multiple insertion sites and the risk of gene disruption. When looking at the genes that are up or down-regulated in CYP2D6^+^/GFP^+^ cells, their chromosomal locations are variable although chromosomes 7, 12, 19 and 22 were found several times. In addition, we did not perform cell cloning after lentiviral infection since HepaRG cells usually do not maintain their ability to fully differentiate after selection of clones. Thus, the CYP2D6^+^/GFP^+^ cell line consists in an heterogenous cell population with multiple different insertions sites probably resulting in various alterations of gene expression. However, we found several genes that were significantly increased in CYP2D6^+^/GFP^+^ cell line such as TRIM63 and NXF3. TRIM63 is an E3 ubiquitin ligase of the muscle tissue and a member of the TRIM protein super-family (Chen et al., 2023; Peris-Moreno et al., 2020). NXF3 is a nuclear export factor used by small viral RNA (Yin et al., 2013). It has also been reported that TRIM family members are antiviral factors induced by multiple viruses including HIV-1 and Hepatitis B virus (Pagani et al., 2021 ; Horvath et al., 2023 ; Shen et al., 2024). Their strong increase in mRNA levels in CYP2D6^+^/GFP^+^ cells may result from a cell adaptation to the expression of the mRNA encoded by the lentiviral transgene, which contains the non-coding WPRE sequence.

## V ACKNOWLEDGEMENTS

The authors would like to thank Stéphanie Dutertre from the Mric microscopy core facility (Federative Research Structure Biosit, Rennes, France) for helping with fluorescence microscopy. The authors would also like to thank Dr. Nicolas Crouvezier, project manager at the licensing department Inserm transfer for his assistance in the writing of the collaboration contract between the laboratory Inserm UMR1317 and the Biotech company Biopredic^International^.

## VI AUTHOR CONTRIBUTIONS

Conceptualization, H.C-E., A.J, C.C., A.C. and P.L.; methodology, H.C-E., A.J, C.B., T.D., T.G., R.P., E.S., and P.L.; validation, A.J, C.C., A.C. and P.L.; formal analysis, H.C-E., A.J, A.C. and P.L.; investigation, H.C-E., F.M., C.B., T.D., C.R., T.G., R.P., F.R., E.S. and A.J. ; data curation, H.C-E., A.J, and P.L.; writing—original draft preparation, H.C-E. and P.L.; writing—review and editing, H.C-E., A.J, C.C., A.C. and P.L. ; supervision, A.J, C.C. and P.L.; project administration, A.J, C.C. and P.L.; funding acquisition, A.J, C.C. and P.L. All authors have read and agreed to the published version of the manuscript.

## VII FUNDING

This work was funded by the Institut National de la Santé et de la Recherche Médicale (Inserm, France). Hugo Coppens-Exandier was a recipient of a fellowship “Convention Industrielle de Formation par la Recherche » (CIFRE n° 221206A10) from the Association Nationale de la Recherche et de la Technologie (ANRT).

## Notes

### Competing Interest Statement

The authors have declared no competing interest.

