## Supplementary material for "Stable lentiviral-mediated expression of Cytochrome P450 2D6 in HepaRG cells: New means for *in vitro* assessment of xenobiotic biotransformation and cytotoxicity": Supporting Information 1.pptx

### Slide 1
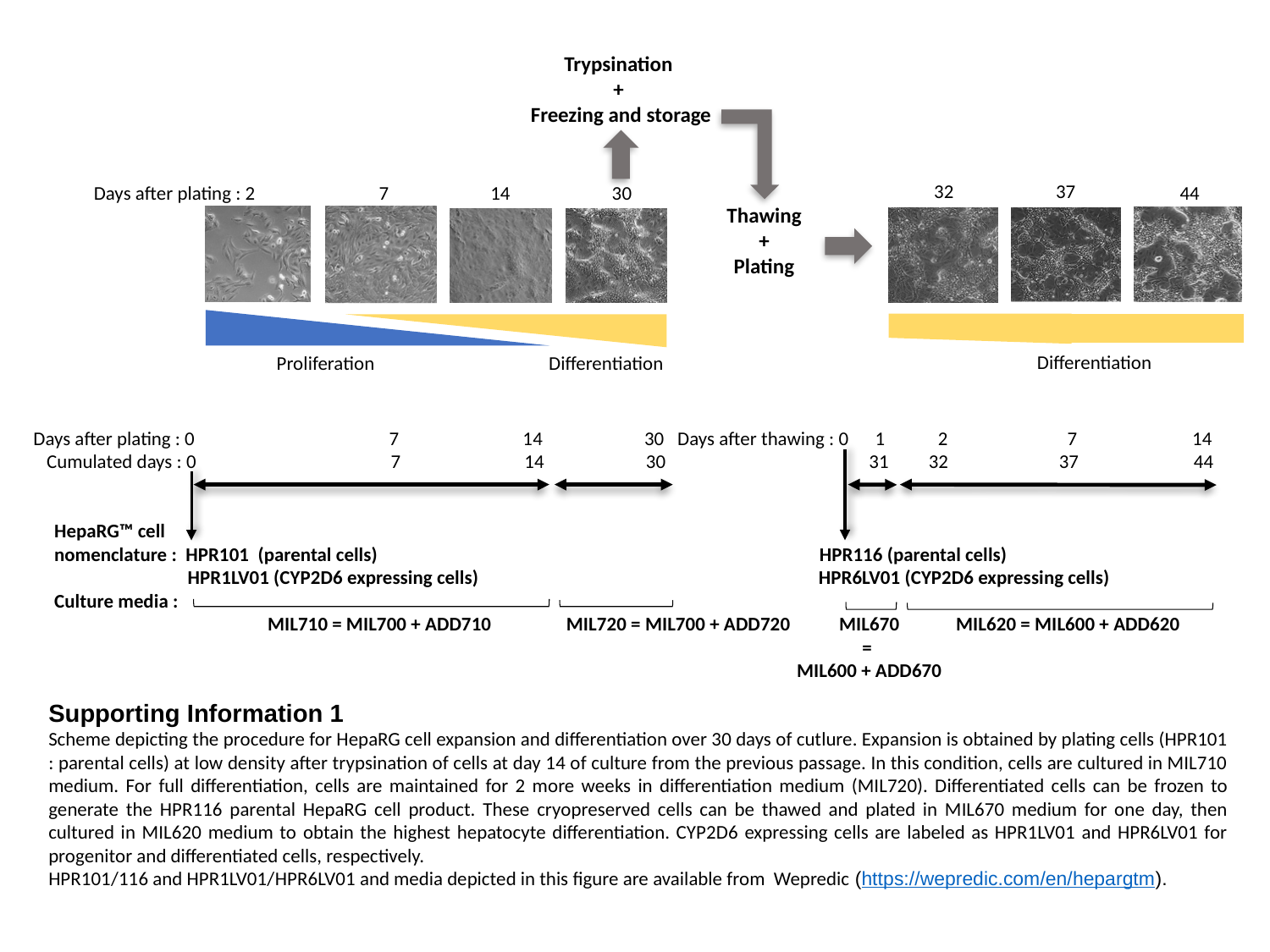

Trypsination
+
Freezing and storage
37
32
Days after plating : 2 7 14 30
44
Thawing
 +
Plating
Differentiation
Differentiation
Proliferation
Days after plating : 0 7 14 30 Days after thawing : 0 1 2 7 14
 Cumulated days : 0 7 14 30 31 32 37 44
HepaRG™ cell
nomenclature : HPR101 (parental cells) HPR116 (parental cells)
 HPR1LV01 (CYP2D6 expressing cells) HPR6LV01 (CYP2D6 expressing cells)
Culture media :
 MIL710 = MIL700 + ADD710 MIL720 = MIL700 + ADD720 MIL620 = MIL600 + ADD620
MIL670
=
MIL600 + ADD670
Supporting Information 1
Scheme depicting the procedure for HepaRG cell expansion and differentiation over 30 days of cutlure. Expansion is obtained by plating cells (HPR101 : parental cells) at low density after trypsination of cells at day 14 of culture from the previous passage. In this condition, cells are cultured in MIL710 medium. For full differentiation, cells are maintained for 2 more weeks in differentiation medium (MIL720). Differentiated cells can be frozen to generate the HPR116 parental HepaRG cell product. These cryopreserved cells can be thawed and plated in MIL670 medium for one day, then cultured in MIL620 medium to obtain the highest hepatocyte differentiation. CYP2D6 expressing cells are labeled as HPR1LV01 and HPR6LV01 for progenitor and differentiated cells, respectively.
HPR101/116 and HPR1LV01/HPR6LV01 and media depicted in this figure are available from Wepredic (https://wepredic.com/en/hepargtm).
