## Supplementary material for "Stable lentiviral-mediated expression of Cytochrome P450 2D6 in HepaRG cells: New means for *in vitro* assessment of xenobiotic biotransformation and cytotoxicity": Supporting Information 2-5.pptx

### Slide 1
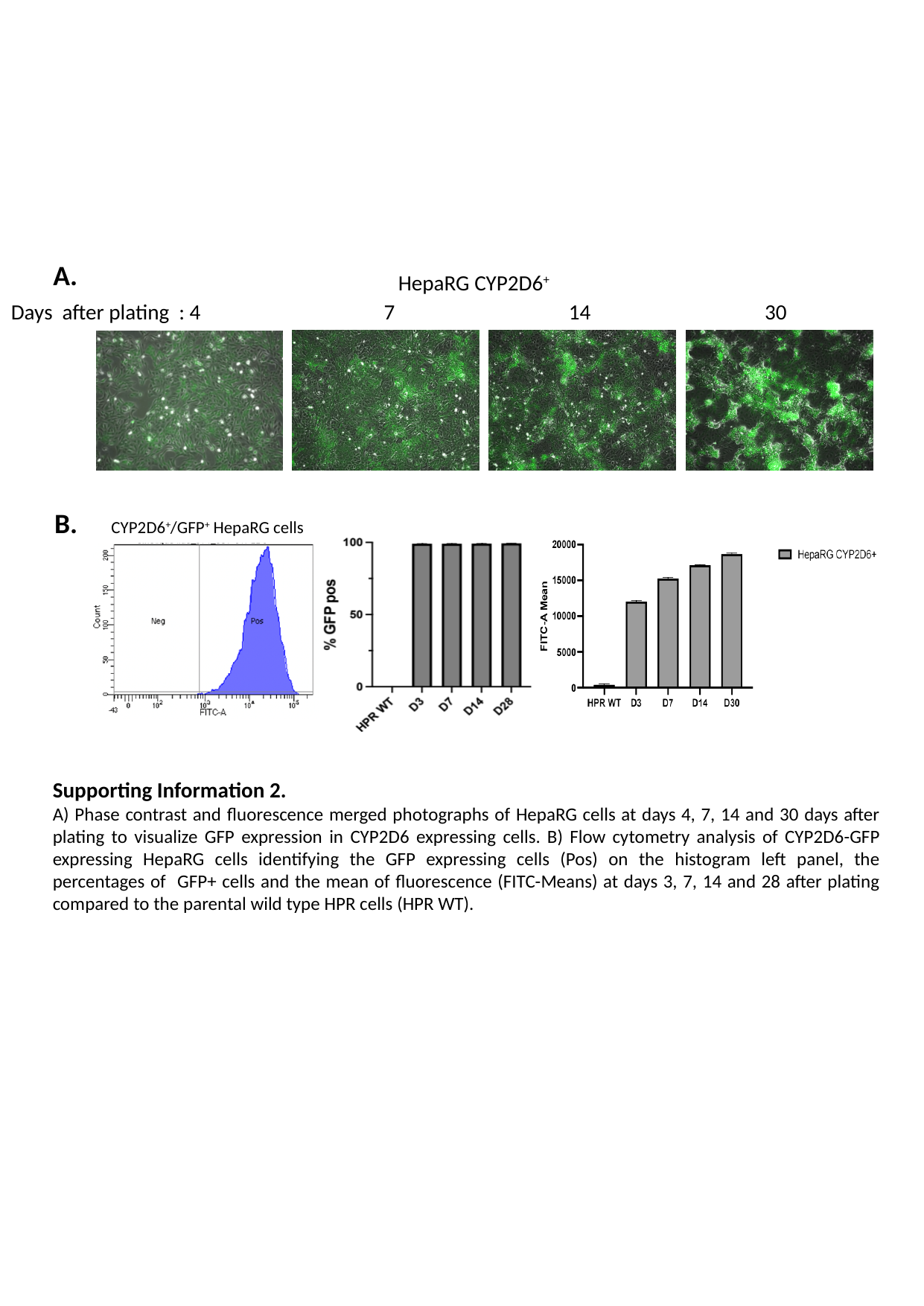

A.
HepaRG CYP2D6^+^
Days after plating : 4 7 14 30
B.
CYP2D6^+^/GFP^+^ HepaRG cells
Supporting Information 2.
A) Phase contrast and fluorescence merged photographs of HepaRG cells at days 4, 7, 14 and 30 days after plating to visualize GFP expression in CYP2D6 expressing cells. B) Flow cytometry analysis of CYP2D6-GFP expressing HepaRG cells identifying the GFP expressing cells (Pos) on the histogram left panel, the percentages of GFP+ cells and the mean of fluorescence (FITC-Means) at days 3, 7, 14 and 28 after plating compared to the parental wild type HPR cells (HPR WT).

### Slide 2
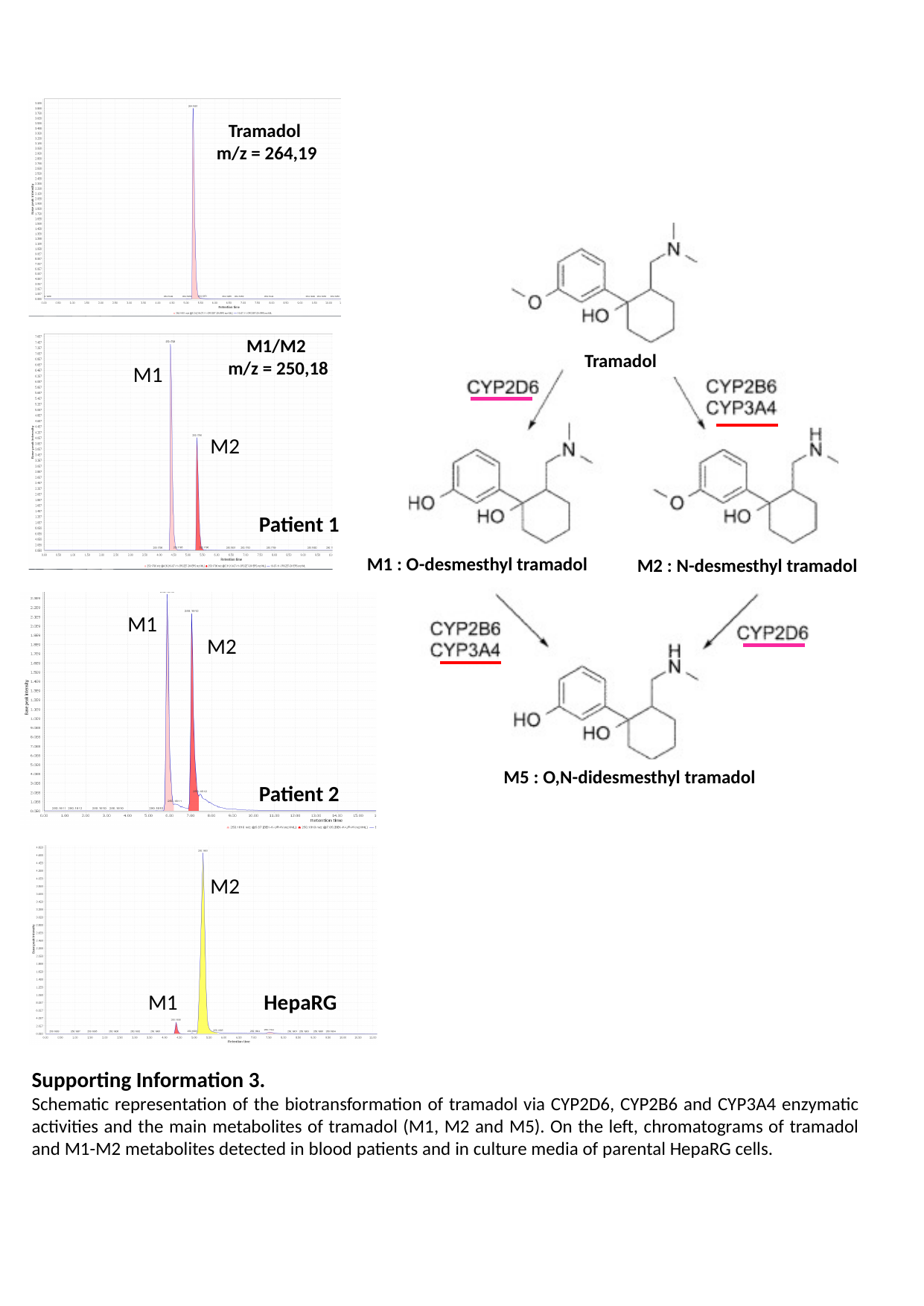

Tramadol
m/z = 264,19
Tramadol
M1 : O-desmesthyl tramadol
M2 : N-desmesthyl tramadol
M5 : O,N-didesmesthyl tramadol
M1/M2
m/z = 250,18
M1
M2
Patient 1
M1
M2
Patient 2
M2
M1
HepaRG
Supporting Information 3.
Schematic representation of the biotransformation of tramadol via CYP2D6, CYP2B6 and CYP3A4 enzymatic activities and the main metabolites of tramadol (M1, M2 and M5). On the left, chromatograms of tramadol and M1-M2 metabolites detected in blood patients and in culture media of parental HepaRG cells.

### Slide 3
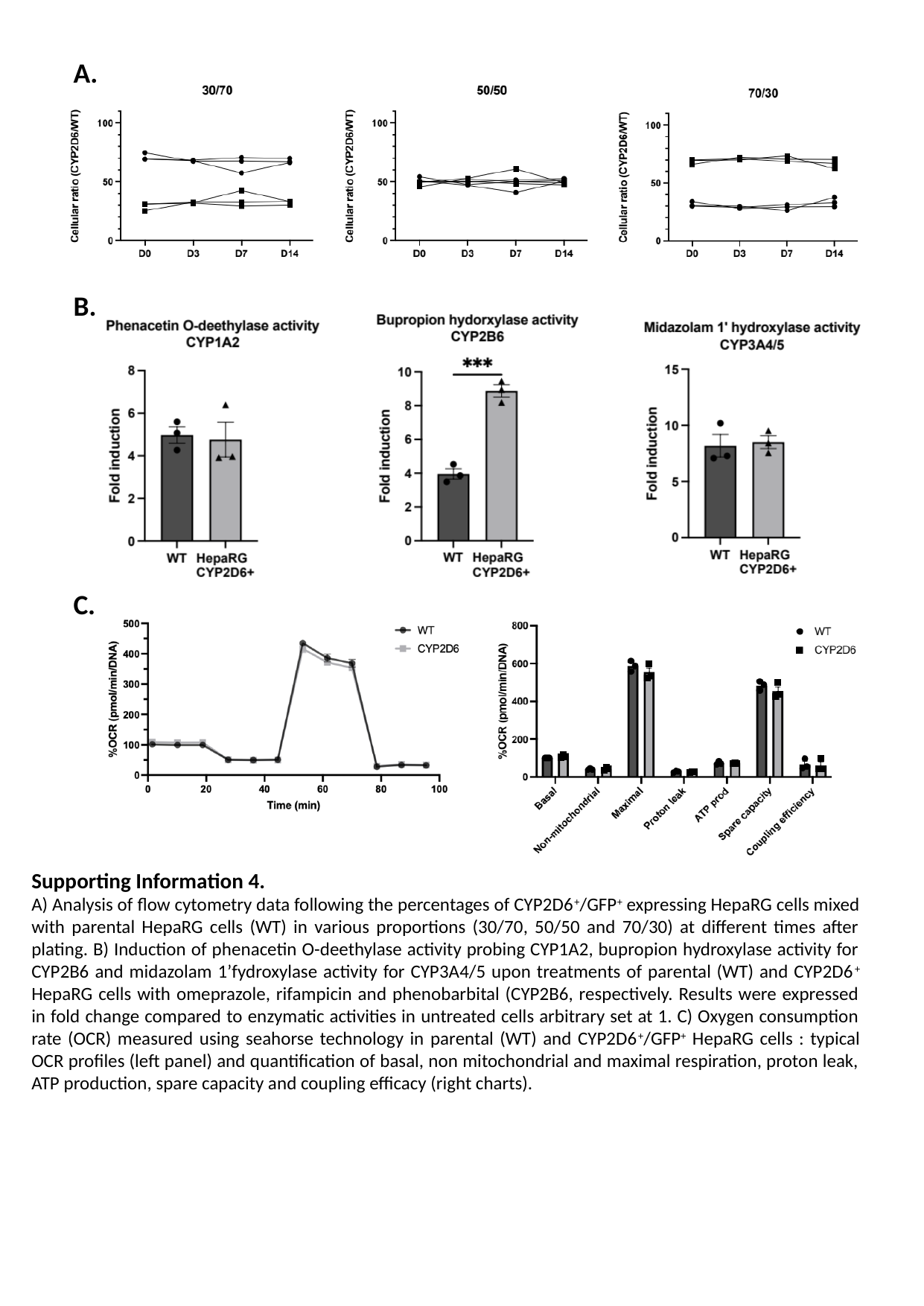

A.
B.
C.
Supporting Information 4.
A) Analysis of flow cytometry data following the percentages of CYP2D6^+^/GFP^+^ expressing HepaRG cells mixed with parental HepaRG cells (WT) in various proportions (30/70, 50/50 and 70/30) at different times after plating. B) Induction of phenacetin O-deethylase activity probing CYP1A2, bupropion hydroxylase activity for CYP2B6 and midazolam 1’fydroxylase activity for CYP3A4/5 upon treatments of parental (WT) and CYP2D6^+^ HepaRG cells with omeprazole, rifampicin and phenobarbital (CYP2B6, respectively. Results were expressed in fold change compared to enzymatic activities in untreated cells arbitrary set at 1. C) Oxygen consumption rate (OCR) measured using seahorse technology in parental (WT) and CYP2D6^+^/GFP^+^ HepaRG cells : typical OCR profiles (left panel) and quantification of basal, non mitochondrial and maximal respiration, proton leak, ATP production, spare capacity and coupling efficacy (right charts).

### Slide 4
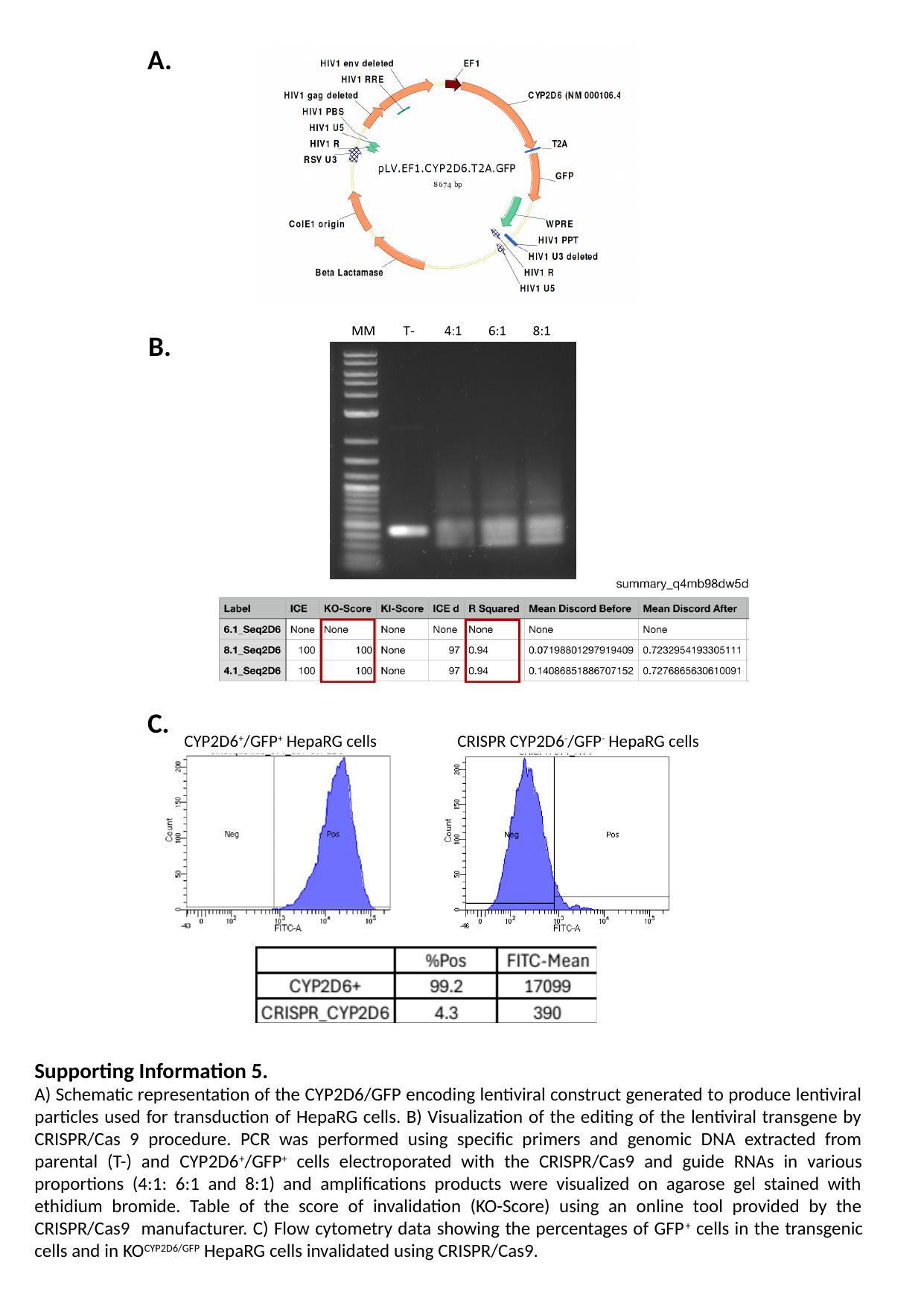

A.
B.
C.
CYP2D6^+^/GFP^+^ HepaRG cells
CRISPR CYP2D6^-^/GFP^-^ HepaRG cells
Supporting Information 5.
A) Schematic representation of the CYP2D6/GFP encoding lentiviral construct generated to produce lentiviral particles used for transduction of HepaRG cells. B) Visualization of the editing of the lentiviral transgene by CRISPR/Cas 9 procedure. PCR was performed using specific primers and genomic DNA extracted from parental (T-) and CYP2D6^+^/GFP^+^ cells electroporated with the CRISPR/Cas9 and guide RNAs in various proportions (4:1: 6:1 and 8:1) and amplifications products were visualized on agarose gel stained with ethidium bromide. Table of the score of invalidation (KO-Score) using an online tool provided by the CRISPR/Cas9 manufacturer. C) Flow cytometry data showing the percentages of GFP^+^ cells in the transgenic cells and in KO^CYP2D6/GFP^ HepaRG cells invalidated using CRISPR/Cas9.
