## Supplementary material for "Stable lentiviral-mediated expression of Cytochrome P450 2D6 in HepaRG cells: New means for *in vitro* assessment of xenobiotic biotransformation and cytotoxicity": Supporting Information Tables 1-2-3.docx

| Guide RNA_#1 | GACUACGGUCAUCACCCACC |
| --- | --- |
| Guide RNA_#2 | CGGCCCGAAACCCAGGAUCU |
| Guide RNA_#3 | CGCGAGGCGCUGGUGACCCA |
| forward_primer | CCACTCGTCACAAGCCCC |
| reverse_primer | GGGGTCGTCCAAGGTTCAAA |
| sequencing_primer | CATGCTCACACCTCCCTAGTG |

**Supporting Information -Table 1**. sgRNA and primer sequences used for CRIPSR-Cas9 experiments

| **Compound** | **Formula** | **RT (min)** | **Precursor ion (*m/z*)** | **Product ion (*m/z*)** | **Declustering potential (V)** | **Collision energy (V)** |
| --- | --- | --- | --- | --- | --- | --- |
| Tramadol | C_16_H_25_NO_2_ | 4.99 | 264.2 | 58.1* | 15 | 7 |
| M1 | C_15_H_23_NO_2_ | 3.85 | 250.2 | 58.0*  232.1 | 30  30 | 22  10 |
| M2 | C_15_H_23_NO_2_ | 5.05 | 250.2 | 44.0*  232.1 | 30  30 | 11  7 |
| M3 | C_14_H_21_NO_2_ | 4.93 | 236.2 | 218.1*  189.1 | 16  16 | 6  12 |
| M4 | C_13_H_16_NO_2_ | 3.85 | 222.1 | 175.1*  204.1 | 18  18 | 12  7 |
| M5 | C_14_H_21_NO_2_ | 3.94 | 236.2 | 44.0*  218.2 | 23  23 | 10  6 |
| Tramadol 13C-D_3_ (IS) | C_16_H_22_D_3_NO_2_ | 4.97 | 268.2 | 58.0* | 30 | 20 |

**Supporting Information-Table 2.** Optimized parameters and MRM transition for quantitation of tramadol, its metabolites and internal standard (IS). *product ion (*m/z*) used for quantitation

Tramadol and its metabolites were measured by liquid chromatography-tandem mass spectrometry (LC-MS/MS) using a Xevo TQ-XS quadrupole mass spectrometer with an ESI ionization source. The instrument was coupled to a Waters Acquity Class-H Plus equipped with a quaternary pump system (Waters, MA, USA). The column was held at 40°C, and the autosampler was set at 15°C. Two microliters of sample were injected, and LC separation was performed on an Acquity UPLC BEH C18 column (2.1 × 100 mm, 1.7 μm; Waters Inc., MA) with a binary gradient of elution at a flow rate of 0.5 mL/min, consisting of ammonium acetate 10 mM/0.1% formic acid (solvent A) and 0.1% formic acid in acetonitrile (solvent B). The gradient started with 5% solvent B for 1.5 min and then ramped up to 30% solvent B over a period of 3.5 min, and up to 40% solvent B over another period of 3 min, and finally up to 95% solvent B within 1 min. The gradient was then maintained at these conditions for 1 min before recycling back to 5% solvent B in 3 min and then held at 5% solvent B for 2 min.

The MS/MS data were acquired in multiple reaction ion monitoring (MRM) mode using positive electrospray ionization. MRM transitions were set up for each analyte: one for quantitation and one for confirmation (as possible); only one MRM transition for tramadol and the IS, as reported in **Table 2**. The following conditions were used: capillary 3 kV, source temperature 150 °C, desolvatation temperature 550 °C, cone gas flow 150 L/h, desolvation gas flow 1000 L/h, and nebulizer 7 bars. Declustering potential and collision energy were optimized to maximize the signal of each reaction monitored.

Seven-point calibration curves were obtained by fortifying drug free human plasma with working solution of tramadol and its metabolites. Standard curves corresponded to peak area ratios of each analytes to IS. Linear concentration ranges from 10 to 1000μg/L for tramadol, from 0.1 to 250μg/L for N,N-bisdesmethyltramadol (M3), from 1 to 500µg/L for O-desmethyltramadol (M1), N-desmethyltramadol (M2), N,O-didesmethyltramadol (M5) and from 0.5 to 250μg/L for O-desmethyl-N,N-bisdesmethyltramadol(M4) were used for quantitation. Tramadol and M1 quality controls (QC) available were analyzed at the beginning and the end of each run.

| CYP3A4-F | CTTCATCCAATGGACTGCATAAA |
| --- | --- |
| CYP3A4-R | TCCCAAGTATAACACTCTACACACACA |
| CYP2D6-F | TAAGGGAACGACACTCATCAC |
| CYP2D6-R | TCACCAGGAAAGCAAAGACAC |
| GFP-F | ACAACAGCCACAACGTCTAT |
| GFP-R | GGGTGTTCTGCTGGTAGTG |
| CYP2A7-F | AGCCCTTGCAGCAACTTAAA |
| CYP2A7-R | CAGTTCACGTCAACCTCAC |
| CYP2C8-F | CATCTGGCTGCCGATCTGCT ATC |
| CYP2C8-R | AGTGACCTGAACAACTCTCCTTAATGG |
| CYP2C9-F | GCCTTTTCTCACCTGTCATCTCAC |
| CYP2C9-R | CAATGCAACTGTTACAGAGTATGGA |
| AhR-F | CTTCAGCCACCATCCATACTT |
| AhR-R | CCTTTGGCATCACAACCAATAG |
| PXR-F | CCAGGACATACACCCCTTTG |
| PXR-R | CTACCTGTGATGCCGAACAA |
| FXR-F | CGACAAGTGACCTCGACAAC |
| FXR-R | GGTCCAAAGTCTGAAATCCTGG |
| HNF4a-F | TCTTTGACCCAGATGCCAAG |
| HNF4a-R | GTCGTTGATGTAGTCCTCCAAG |
| TBP-F | GAGCTGTGATGTGAAGTTTCC |
| TBP-R | TCTGGGTTTGATCATTCTGTA |
| WPRE-F | GTCAGCTCCTTTCCGGGACT |
| WPRE-R | CACCACGGAATTGTCAGTGC |
| CAR-F | ATGCTGGCATGAGGAAAGAC |
| CAR-R | GTTGCACAGGTGTTTGCTGT |
| NXF3-F | AGAAGAGCAAGATGTTGGGATATT |
| NXF3-R | GCTGCTGATGGGATGAAGAA |
| TRIM63-F | CTTCCAGGCTGCAAATCCCTA |
| TRIM63-R | ACACTCCGTGACGATCCATGA |

**Supporting information-Table 3.** Forward (F) and reverse (R) primer sequences.
